# Distal-less homeobox 5 gives rise to myofibroblastic carcinoma-associated fibroblasts to promote collective breast cancer invasion

**DOI:** 10.64898/2026.08.30.745418

**Authors:** Yoshihiro Mezawa, Kohei Kumegawa, Katsuki Morita, Liying Yang, Karin Hirakuri, Kazunari Yamashita, Takuya Shirakihara, Ritsuko Sasaki, Hiroko Onagi, Goro Kutomi, Reo Maruyama, Akira Orimo

**Affiliations:** Department of Pathology and Oncology, Juntendo University Faculty of Medicine, Tokyo, Japan; Department of Molecular Pathogenesis, Juntendo University Graduate School of Medicine, Tokyo, Japan; Cancer Cell Diversity Project, NEXT-Ganken Program, Japanese Foundation for Cancer Research, Tokyo, Japan; Division of Cancer Epigenomics, Cancer Institute, Japanese Foundation for Cancer Research, Tokyo, Japan; Department of Breast Oncology, Juntendo University Graduate School of Medicine, Tokyo, Japan; Department of Breast Surgery, Institute of Science Tokyo, Tokyo, Japan; Department of Human Pathology, Juntendo University School of Medicine, Tokyo, Japan; Faculty of Health Sciences, Graduate School of Medicine, Juntendo University, Tokyo, Japan

## Abstract

Tumor-promoting myofibroblastic carcinoma-associated fibroblasts (myCAFs) are induced by activation of transforming growth factor-β (TGF-β) signaling. However, the molecular basis of myCAF-specific transcriptional programs regulated by TGF-β signaling remains poorly understood. Using a meta-analysis of single-cell RNA-seq data from 132 human breast tumor and non-tumor tissues, we show that myCAFs activate gene regulatory programs relevant to skeletal and cardiovascular development that are associated with poorer outcomes in breast cancer patients. Of note, distal-less homeobox 5 (DLX5), a master transcription factor for skeletal development, is activated in human breast myCAFs at both epigenetic and transcriptional levels. DLX5 expression is also initiated by TGF-β1 treatment in human mammary fibroblasts. Immunoprecipitation and CUT&RUN assays using DLX5-expressing fibroblasts demonstrate that DLX5 interacts with Smad2/3/4 proteins, enabling their cooperative occupancy at shared genomic binding sites of target genes, thereby promoting canonical TGF-β signaling and the myCAF state. DLX5-primed myCAFs also enhance paracrine TGF-β signaling and neuropilin-2 expression to promote collective tumor invasion. Our findings indicate that DLX5 induces myCAF formation and promotes breast tumor progression in collaboration with canonical TGF-β signaling.

## Introduction

Carcinoma-associated fibroblasts (CAFs) are abundant in the tumor-associated stroma of various human carcinomas including those of the breast. Many different cells of origin and various stimuli from the tumor microenvironment (TME) generate heterogeneous CAF subpopulations including myofibroblastic CAFs (myCAFs), inflammatory CAFs (iCAFs) and antigen-presenting CAFs (apCAFs) that have distinct impacts on tumor progression (Bartoschek *et al*, 2018; Elyada *et al*, 2019; Gao *et al*, 2024; Lavie *et al*, 2022; Liu *et al*, 2026; Mezawa & Orimo, 2022).

myCAFs are enriched in the TME and have the ability to remodel and contract the extracellular matrix (ECM), thereby promoting tumor invasion, metastasis and immune exclusion (Krishnamurty *et al*, 2022; Lavie *et al*., 2022; Liu *et al*., 2026; Mariathasan *et al*, 2018). myCAFs also exhibit characteristics resembling those of myofibroblasts, a specialized fibroblast population involved in wound healing and fibrosis (Polanska *et al*, 2010; Serini & Gabbiani, 1999). Accordingly, these CAFs are referred to as myCAFs.

Transforming growth factor-β (TGF-β) produced by myCAFs acts on their TGF-β receptors in an autocrine fashion to establish the myCAF traits (Kojima *et al*, 2010; Massague & Sheppard, 2023). The paracrine TGF-β signaling from CAFs also induces partial epithelial-mesenchymal transition and an immunosuppressive milieu in tumors (Matsumura *et al*, 2019; Nixon *et al*, 2023; Okubo *et al*, 2025). In response to activation of TGF-β, TGF-β receptor type I (TGFBRI/ALK5) is recruited and phosphorylated to stimulate phosphorylation of receptor-regulated Smads (R-Smads) including Smad2 and Smad3 via activation of its kinase activity (David & Massague, 2018). R-Smads then form a complex with Smad4 and physically interact with other transcription factors (TFs), so-called Smad cofactors or lineage-determining transcription factors, to acquire the specificity and affinity of DNA binding sites in various target genes depending on the biological context (David & Massague, 2018; Miyazawa *et al*, 2024). However, cofactors for the R-Smad complex that orchestrate the myCAF state have not been identified.

Distal-less homeobox 5 (DLX5) is a homeobox TF relevant to skeletal development (Acampora *et al*, 1999; Levi *et al*, 2022). DLX5 mutant mice exhibit perinatal lethality and abnormalities in craniofacial development, vestibular organ morphogenesis and osteogenesis (Acampora *et al*., 1999; Depew *et al*, 1999). However, roles of DLX5 associated with skeletal development in CAFs remain unknown.

In this study, we show that a skeletal developmental program regulated by DLX5 is activated in myCAFs in human breast cancers. DLX5 expression is also initiated by TGF-β1 treatment in human mammary fibroblasts, and DLX5 interacts with an R-Smad complex to occupy shared genomic binding sites of target genes, thereby promoting canonical TGF-β signaling and the myCAF state. DLX5-primed myCAFs, in turn, promote breast tumor protrusion via induction of paracrine TGF-β signaling and neuropilin-2 expression.

## Results

### Establishment of integrated single-cell (sc)RNA-seq data of human breast tissues

To reveal microenvironmental changes between human breast cancer and non-cancer tissues, we generated a scRNA-seq dataset by integrating seven publicly available datasets (Bhat-Nakshatri *et al*, 2021; Delgado *et al*, 2024; Gray *et al*, 2022; Murrow *et al*, 2022; Pal *et al*, 2021; Tietscher *et al*, 2023; Wu *et al*, 2021) (Fig. 1A, EV1A, Supplementary Table 1). After quality control and integration, we obtained 371,040 and 173,753 cells from cancerous and noncancerous breasts, respectively. Unsupervised clustering identified 38 clusters, whose cell types were assigned according to expression levels of the lineage markers for immune cells, epithelial cells, endothelial cells (ECs), perivascular cells (PVs: pericytes and smooth muscle cells (SMCs)) and fibroblasts (Fig. 1B, EV1B-E). We notably found the connection between cluster 12 (fibroblasts) and cluster 9 (PVs) on a uniform manifold approximation and projection (UMAP) plot, indicative of a pericyte-to-fibroblast transition (PFT) (an arrowhead in Fig. 1B), consistent with a previous report (Hosaka *et al*, 2016). In contrast, we did not identify any connections between fibroblasts and ECs, a potential source of CAFs in mice (Zeisberg *et al*, 2007). Taken together, these findings suggest that a particular human breast fibroblast population shares characteristics of PVs.

**Figure 1.**
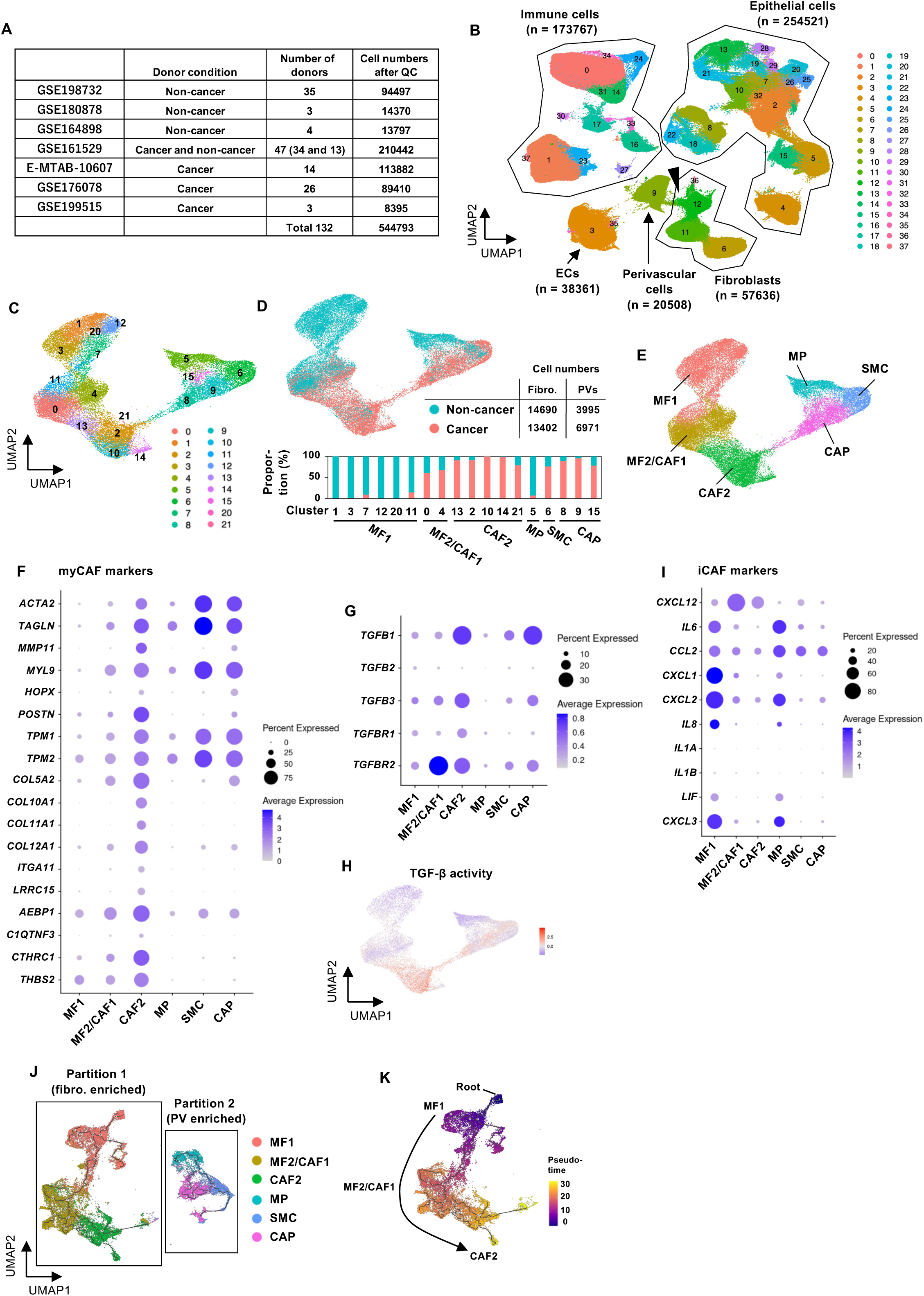
Heterogeneity of fibroblasts and PVs in human breast tumor and non-tumor tissues. **(A)** Information about the sources of the data which we used for the integration. QC, quality control. **(B)** Uniform manifold approximation and projection (UMAP) of all 544,793 cells colored by clusters. An arrowhead highlights a connection between fibroblasts and perivascular cells. **(C-E)** UMAP of 39,058 fibroblasts and perivascular cells (PVs), which are colored by cluster ID (C), donor condition (D) and cluster type (E). MF, mammary fibroblasts; MP, mammary pericytes; CAP, carcinoma-associated pericytes; SMC, smooth muscle cells. **(F, G)** Balloon plots of myCAF markers and TGF-β ligands and receptors. **(H)** Activity of TGF-β signaling using decoupleR tools. **(I)** Balloon plots of iCAF markers. **(J)** Dimensionality reduction plot generated by Monocle3 is shown with the cluster types defined in (E). **(K)** Pseudotime was visualized on the UMAP shown in (J). Lines on cells indicate the exact trajectory, whereas the curved arrow is drawn to aid interpretation of the data.

### Identification of six major clusters among fibroblasts and PVs

To investigate precisely gene expression programs of fibroblasts and PVs, these cells were subjected to subclustering. After removal of possible doublets and uncharacterized clusters from the initial 23 clusters (Fig. EV1F), we obtained 18 clusters including 39,058 cells (Fig. 1C, D). According to enrichment of cells expressing markers for fibroblasts (*decorin* (*DCN*) and *platelet-derived growth factor receptor alpha* (*PDGFRA*)) and PVs (*myosin heavy chain 11* (*MYH11*) and *calponin 1* (*CNN1*) for SMCs, and *regulator of G protein signaling 5* (*RGS5*) for pericytes), we considered the left side as fibroblasts and the right side as PVs (Kumar *et al*, 2023) (Fig. 1C, EV1G). Fibroblasts in clusters 1, 3, 7, 12, 20 and 11 were derived from non-cancerous donors, while those in clusters 13, 2, 10, 14 and 21 were derived from cancer patients (Fig. 1D). Clusters 0 and 4 were intermediate (non-cancerous 60.5% and 66.4%; cancerous 39.5% and 33.6%, respectively) (Fig. 1D). Based on these results, we summarized and named these clusters as mammary fibro 1 (MF1), MF2/CAF1 mix and CAF2 (Fig. 1D, E).

Next, according to the distribution of a pericyte marker and SMC markers, we considered clusters 5, 8, 9 and 15 as pericytes and cluster 6 as SMCs (Fig. 1E, EV1G). Although pericytes in cluster 8 expressed fibrotic genes (*collagen type I alpha 1 chain* (*COL1A1*) and *fibronectin 1* (*FN1*)), we regarded them as cells undergoing PFT rather than a fibroblast subset because of their low *PDGFRA* expression (Fig. EV1G, H). Collectively, fibroblasts and PVs were grouped into six major clusters.

### myCAFs are enriched in the CAF2 cluster and arise from fibroblasts in an iCAF state

CAFs comprise myCAF, iCAF, and apCAF subtypes, which exhibit substantial plasticity (Croizer *et al*, 2024; Krishnamurty *et al*., 2022; Wang *et al*, 2023). To investigate the six major fibroblast and PV clusters, we plotted them using markers for major CAF subtypes (Dominguez *et al*, 2019; Elyada *et al*., 2019). Expression levels of most myCAF markers (Fig. 1F), TGF-β ligands, their receptors (Fig. 1G) and TGF-β activity (Fig. 1H) were progressively increased from MF1 to CAF2, consistent with our previous report that indicated the establishment of TGF-β-autocrine signaling in myCAFs (Kojima *et al*., 2010). In contrast, expression levels of iCAF markers were gradually decreased from MF1 to CAF2 excepting *C-X-C motif chemokine ligand 12* (*CXCL12*) (Fig. 1I). MF1 with the increased iCAF markers was consistently demonstrated as a subset of mammary fibroblasts, named “fibro-major,” which expresses *CXCL1* and *CXCL2* (Kumar *et al*., 2023). apCAF markers were sparsely distributed across the CAF2 cluster, with the highest expression observed in cluster 21 (Fig. EV1I, J). These data indicate myCAFs to be enriched in the CAF2 cluster relative to the CAF1 and MF1 clusters.

To examine the cellular origins of myCAFs, pseudotime trajectory analysis was performed using Monocle3. Fibroblasts and PVs were separated into two partitions according to the clustering algorithm (Fig. 1J). Importantly, we found a trajectory progressing from MF1 through MF2/CAF1 to the CAF2 cluster (Fig. 1K), suggesting a potential transition from an iCAF state to a myCAF state in human breast cancers.

### Enrichment of cardiovascular and skeletal development genes in the CAF2 cluster

To characterize the clusters globally, positive marker genes for each cluster were identified using statistical tests and subsequently used as gene signatures in downstream analyses. The MF2/CAF1 and CAF2 gene signatures comprised 98 and 252 genes, respectively (Fig. 2A, Supplementary Table 2). Gene Ontology (GO) analysis revealed no consistently enriched biological processes in the CAF1 gene signature (Fig. 2B). In contrast, the CAF2 gene signature was enriched for “collagen fibril organization,” “regulation of TGF-β receptor signaling pathway,” and multiple GO terms related to cardiovascular and skeletal system development (Fig. 2C). Consistent with these findings, the CAF2 signature was associated with various cardiovascular and skeletal abnormalities and diseases based on analyses using the Human Phenotype Ontology and Mouse Genome Database implemented in PhenoExam (Cisterna et al., 2022) (Fig. EV2A).

**Figure 2.**
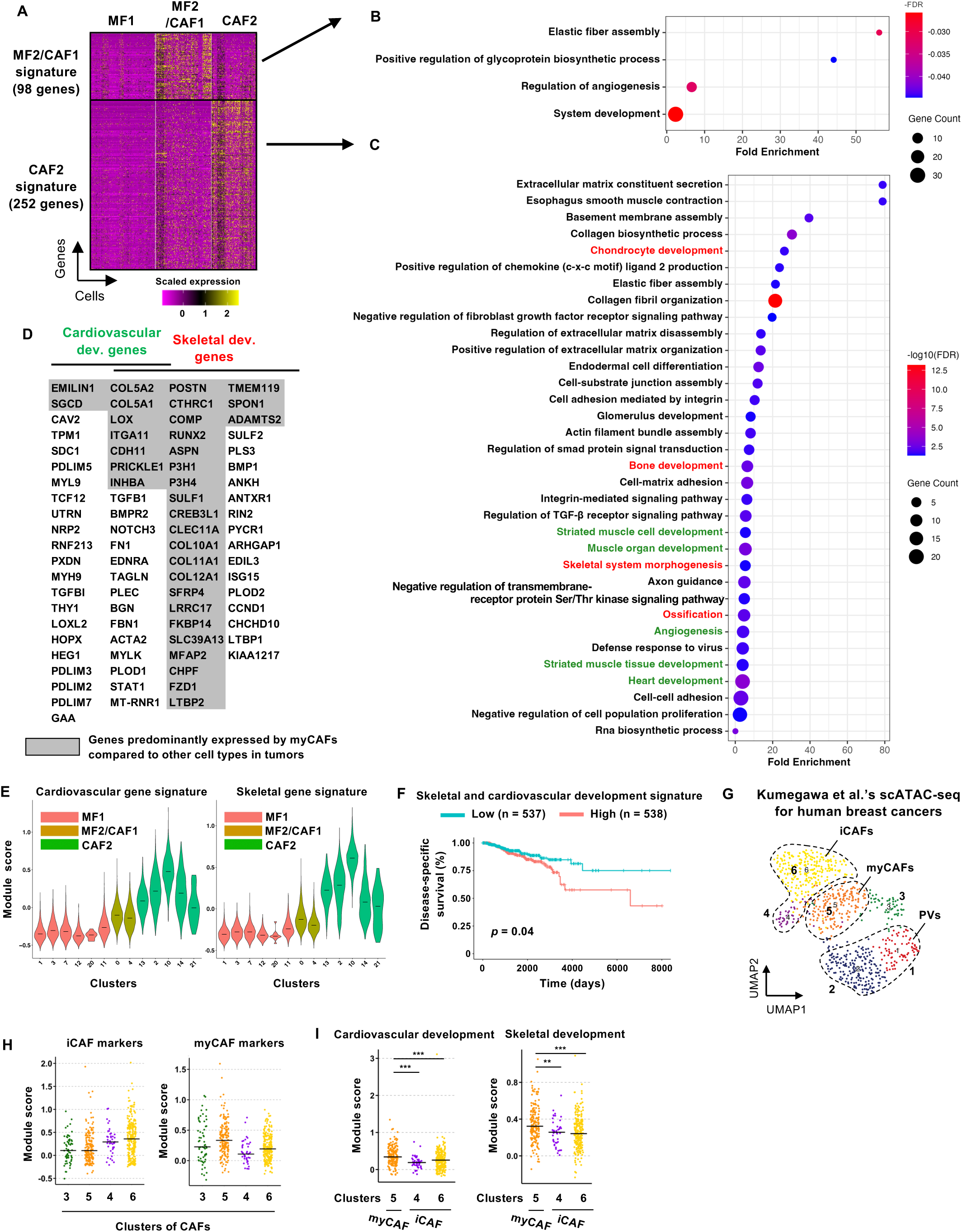
The biological characteristics of human breast CAFs. **(A)** Positive markers selected by the Mann-Whitney U test for MF2/CAF1 or CAF2 were shown as a heatmap. **(B, C)** GO terms meeting FDR < 0.05 in the positive markers of either MF2/CAF1 or CAF2. *p*-values were determined by Fisher’s exact test. GO terms associated with the cardiovascular or skeletal system are shown in green and red, respectively. **(D)** Genes related to GO terms associated with the cardiovascular or skeletal system in (C) or to phenotypic terms in Figure EV2A were listed. **(E)** Scores for the genes specific to either cardiovascular or skeletal development in (D) were calculated. Clusters correspond to Figure 1C. Horizontal lines, median. **(F)** Kaplan-Meier plot of breast cancer data obtained from TCGA. The cardiovascular and skeletal developmental signatures were defined as genes predominantly expressed by myCAFs (highlighted by gray background in (D)), to enable evaluation of effects associated with CAFs. Patients were split into two groups based on the median value. *p*-value was determined by the Log-Rank test. **(G)** UMAP based on scATAC-seq data of human breast cancers. **(H, I)** Module score analysis of the scATAC-seq data. Module scores based on gene activity scores for the gene lists shown in Fig. 1F and I are shown in (H). Those based on the genes specific to either cardiovascular or skeletal development in (D) are shown in (I). Dots, cells; horizontal bars, medians. ** *p* < 0.01 and *** *p* < 0.001 by the Mann-Whitney U test with Benjamini-Hochberg correction (n = 43, 182 and 242 for clusters 4, 5 and 6, respectively).

Genes related to the cardiovascular and skeletal developmental programs accounted for 17.1% (43 genes) and 23.8% (60 genes), respectively, in the CAF2 cluster (252 genes) and 21 genes were shared between the cardiovascular and skeletal developmental genes (Fig. 2D). 33 of these 82 genes were predominantly expressed in myCAFs (Fig. 2D, EV2B).

We found the highest cardiovascular and skeletal development scores in cluster 10 of CAF2 (Fig. 2E) that includes tumor-promoting myCAF markers, such as *leucine-rich repeat-containing protein 15* (*LRRC15*) (Krishnamurty *et al*., 2022), *integrin subunit alpha 11* (*ITGA11*) (Zheng *et al*, 2024) and *cyclin-dependent kinase inhibitor 2A* (*CDKN2A*), which encodes p16, a marker of senescent CAFs (Ye *et al*, 2024) (Fig. EV2C). A high level of the cardiovascular and skeletal developmental signature, defined by genes predominantly expressed in myCAFs (Fig. EV2B), was also associated with poorer disease-specific survival in breast cancer patients (Fig. 2F). Collectively, these findings suggest that increased cardiovascular and skeletal developmental programs might mediate the tumor-promoting ability in myCAFs.

To investigate whether modulation of chromatin accessibility underlies activation of the cardiovascular and skeletal developmental programs in myCAFs, we performed bioinformatic analysis on in-house-published scATAC-seq data from 16 human breast cancer patients (Kumegawa *et al*, 2022). The gene activity score (GS), which is globally correlated with mRNA expression (Granja *et al*, 2021), was calculated based on these data. Based on the GSs of major lineage markers, 527 CAFs and 277 PVs were isolated from a total of 13,788 cells derived from human breast cancers (Fig. EV3A-C). The subclustering and module score analyses identified groups corresponding to myCAFs (cluster 5), iCAFs (clusters 4, 6) and PVs (clusters 1, 2) (Fig. 2G, H, EV3C, D). Cluster 3 could not be clearly assigned to any of these groups, although it showed a trend toward a myCAF state (Figs. 2G, H, EV3C, D). Importantly, we found significantly higher module scores generated by the cardiovascular and skeletal developmental genes in myCAFs relative to iCAFs (Fig. 2I). Taken together, these findings indicate that the cardiovascular and skeletal developmental signatures are epigenetically reprogrammed in myCAFs in breast cancer patients.

### Identification of TFs regulating the cardiovascular and skeletal developmental programs in myCAFs

Given the significance of the cardiovascular and skeletal developmental programs in myCAFs in breast cancer patients, we sought to identify key TFs regulating these programs using TF activity prediction from the scRNA-seq data. We found activation of DLX5 and runt-related transcription factor 2 (RUNX2) regulating skeletal development (Acampora *et al*., 1999; Komori *et al*, 1997) and conversely deactivation of msh homeobox 2 (MSX2), a repressor of RUNX2 (Shirakabe *et al*, 2001), in myCAFs (Fig. 3A, B). TFs associated with cardiovascular development were not among the top 20 most variable TFs in CAF2.

**Figure 3.**
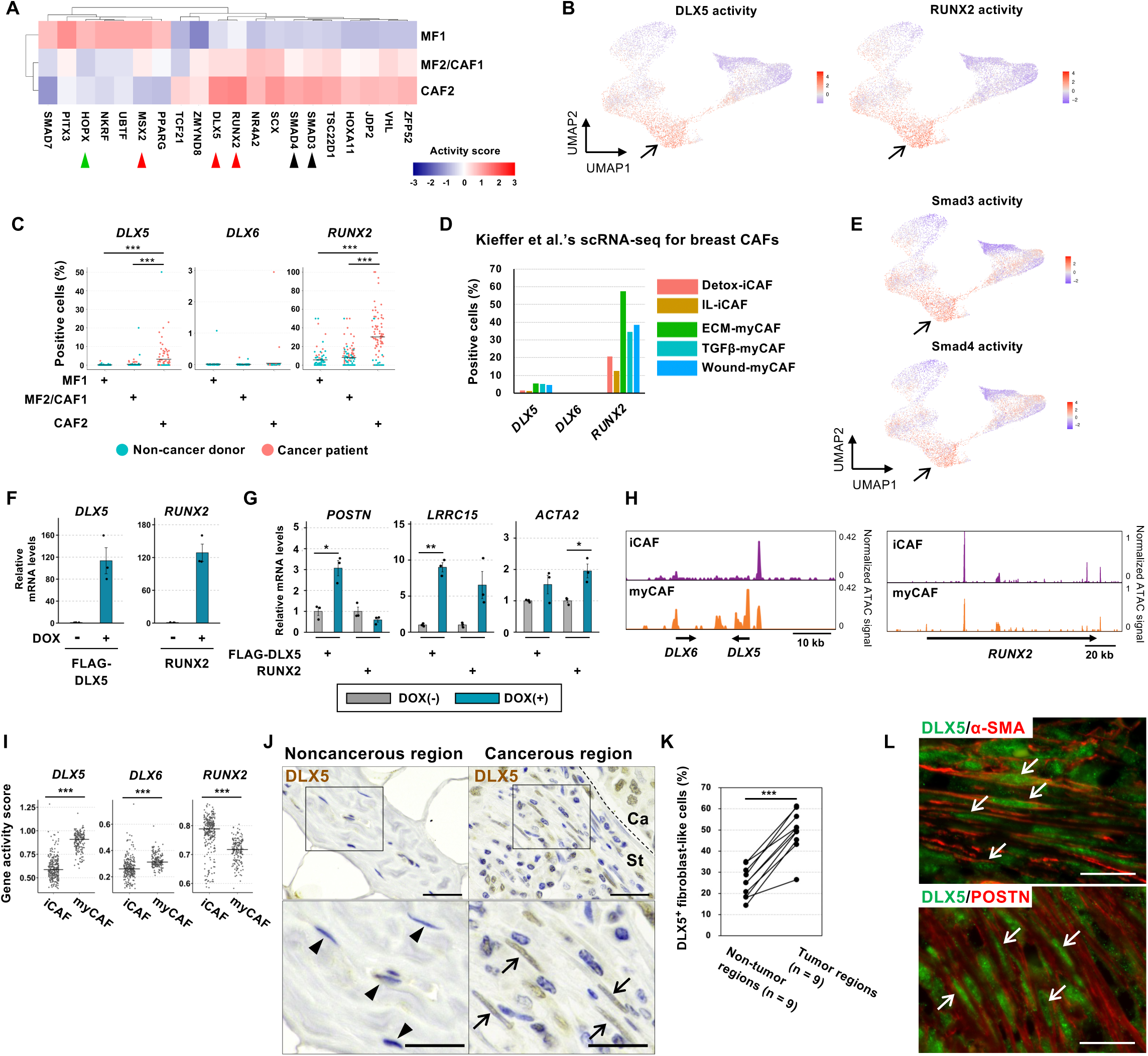
Increased DLX5 activity and expression in myCAFs of human breast cancers. **(A)** Transcription factors’ activity was estimated using the scRNA-seq data. Top 20 TFs showing the greatest variability in mean activity scores across fibroblast clusters are shown. DLX6 was not included in the database used here. Green and red arrowheads, cardiovascular and skeletal developmental TFs, respectively. Black arrowheads, TGF-β signaling. **(B)** Activity scores for each cell in (A) are shown as a heatmap. Arrows, the CAF2 cluster. **(C)** For each patient, the proportion of cells expressing the indicated TFs was calculated. **(D)** Proportions of cells expressing indicated TFs in scRNA-seq data generated by Dr. Kieffer and colleagues (Kieffer *et al*., 2020). **(E)** Activity scores for each cell in (A) are shown as a heatmap. Arrows, the CAF2 cluster. **(F, G)** Real-time PCR for HMFs introduced with doxycycline (DOX)-inducible lentiviral vectors harboring indicated TFs. RNA was extracted at day 4 from DOX induction (0.5 µg/ml). \**p* < 0.05, **p < 0.01 by Welch’s t-test (n = 3). Error bars, standard error (SE). **(H)** Pseudobulk clusters were generated using cells of cluster 5 (myCAFs) and clusters 4 and 6 (iCAFs) in Figure 2G. **(I)** Gene scores are shown for myCAFs and iCAFs in the scATAC-seq data of human breast cancers. \*\*\**p* < 0.001 by the Mann-Whitney U test. **(J, K)** Immunohistochemistry for DLX5 in human breast cancer tissues. \*\*\**p* < 0.001 by the paired t-test (n = 9). Ca, cancer; St, stroma. Arrows and arrowheads indicate DLX5-positive and -negative cells, respectively. Scales, 30 µm (upper) and 20 µm (bottom). **(L)** Double immunofluorescence staining for DLX5 and α-SMA or POSTN in breast cancer tissue negative for estrogen receptor and progesterone receptor and positive for human epidermal growth factor receptor 2. Arrows, double-positive cells. Scales, 20 µm.

We therefore decided to investigate TFs regulating skeletal development in myCAFs. We found DLX5- and RUNX2-expressing cell proportions to be significantly increased in the CAF2 cluster compared to MF1 and MF2/CAF1 clusters in the integrated dataset (Fig. 3C). In contrast, DLX6, a homolog of DLX5, was hardly detected in myCAFs (Fig. 3C). Consistent with these findings, an independent scRNA-seq dataset of breast CAF-S1 cells also showed increased proportions of DLX5- and RUNX2-expressing cells and elevated DLX5 and RUNX2 activities in myCAFs (Kieffer *et al*, 2020) (Fig. 3D, EV4A). Smad3 and Smad4, essential mediators of TGF-β-Smad signaling, were activated in the CAF2 cluster (Fig. 3A, E), consistent with our earlier findings (Fig. 1G, H). Meanwhile, BMP-Smad1/5/9 signaling, which is also crucial for mesenchymal tissue development, was not exclusively activated in CAF2 (Fig. EV4B). Collectively, these findings indicate increased DLX5, RUNX2 and Smad3/4 activity in myCAFs.

### DLX5 induces myCAF markers in HMFs

Since DLX5 and RUNX2 activity is increased in myCAFs, we reasoned that these TFs may contribute to inducing myCAFs state in fibroblasts. To address this issue, we introduced each of doxycycline (DOX)-inducible cDNA expression constructs into 218TGpp cells, immortalized human mammary fibroblasts (HMFs) (Kojima *et al*., 2010) that strongly induced *DLX5* and *RUNX2* mRNA expression (Fig. 3F). Expression levels of myCAF marker genes, including *periostin* (*POSTN)* and *LRRC15*, were significantly increased in the DOX-treated FLAG-tagged DLX5-expressing HMFs (DLX5-HMFs), compared to the control cells without DOX treatment (Fig. 3G). A trend toward increased expression of *ACTA2* mRNA, which encodes α-smooth muscle actin (SMA), another myCAF marker, was also observed in these cells (Fig. 3G). In contrast, expression of *ACTA2*, but not *POSTN* and *LRRC15* was significantly induced in RUNX2-expressing HMFs. These findings suggest that DLX5 induces the myCAF state in HMFs more strongly than RUNX2 does.

We also assessed chromatin accessibility at the *DLX5* and *RUNX2* loci in CAFs by pseudobulk analysis and GS of scATAC-seq data from human breast cancers. ATAC signal and GS at the *DLX5/6* locus were significantly increased in myCAFs compared with those in iCAFs, and these increases were more intense in the locus at *DLX5* than that at *DLX6* (Fig. 3H, I). These findings suggest that the increased transcriptional activity of DLX5 is associated with increased chromatin accessibility. In contrast, there were no increased ATAC signals at the *RUNX2* locus (Fig. 3H) with increased GS in iCAFs (Fig. 3I), indicating *RUNX2* upregulation in myCAFs to be independent of chromatin accessibility. We therefore prioritized further investigation of the role of DLX5 because its marked upregulation was associated with epigenetic remodeling at myCAF marker genes.

### DLX5 is present in myCAFs of human breast cancers

We next investigated whether DLX5-positive CAFs are present in human breast cancers by immunostaining using an anti-DLX5 antibody (see Methods for subtype information). Stronger DLX5 staining was detected in fibroblast-like cells on tumor tissues compared with those on adjacent noncancerous stroma in the same patients (Fig. 3J). DLX5 staining was also detected in breast cancer cells (Fig. 3J), consistent with a previous report (Morini *et al*, 2010). In nine breast cancer patients, we found significantly higher DLX5-positive fibroblast-like cell proportions in cancer regions (49.4%) relative to noncancerous regions (25.1%) (Fig. 3K). DLX5 staining was detected in α-SMA- and POSTN-positive myCAFs in tumor-associated stroma (Fig. 3L), suggesting that DLX5 is expressed in myCAFs in human breast cancers.

### DLX5 cooperates with canonical TGF-β signaling to promote the myCAF state

Since myCAF marker genes were upregulated in HMFs by DLX5, we examined TGF-β signaling in DLX5-HMFs by western blotting. Levels of phosphorylated Smad2 (pSmad2), POSTN and LRRC15 proteins were substantially increased compared to those in the control GFP-HMFs (Fig. 4A). Treatment with SB431542, an inhibitor of TGFBRI, strongly attenuated the elevated pSmad2 protein (Fig. 4B), *POSTN* and *LRRC15* mRNA (Fig. 4C) in DLX5-HMFs, indicating the DLX5-induced myCAF state to be mediated by TGF-β-Smad2/3 signaling in these fibroblasts.

**Figure 4.**
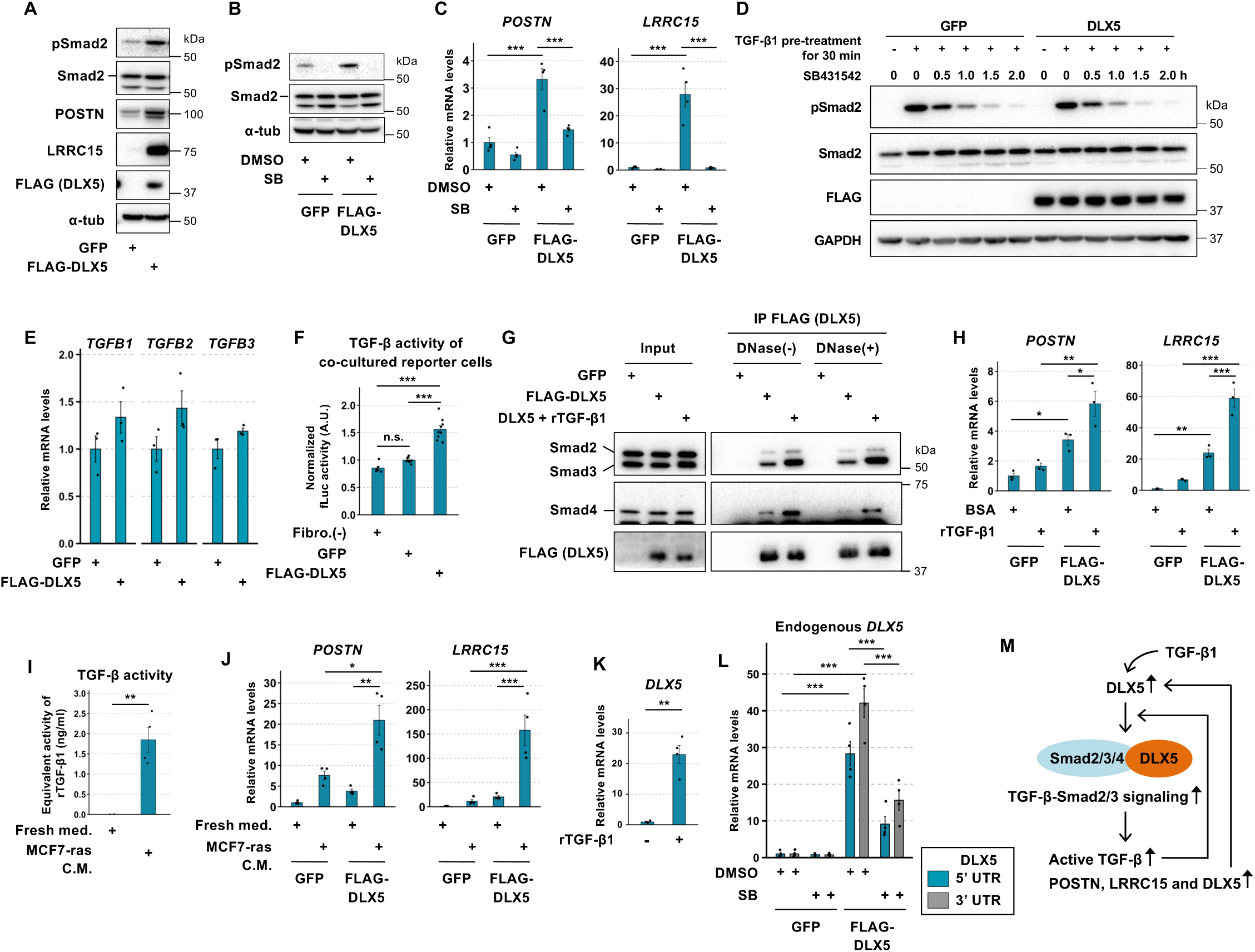
DLX5 activates canonical TGF-β signaling via interaction with Smad2/3/4 in HMFs. **(A)** Western blotting (WB) for HMFs introduced with DOX-inducible vectors for indicated genes (GFP- or DLX5-HMFs). Those cells were treated with DOX for 4 days (0.5 µg/ml). **(B, C)** WB and real-time PCR for GFP- or DLX5-HMFs, which were treated with SB431542 (2 µM) and DOX (0.5 µg/ml) for 4 days. *p*-values were determined by Tukey’s test (n = 4) in (C). **(D)** GFP- or DLX5-HMFs treated with DOX (0.5 µg/ml) for 48 h were stimulated with recombinant TGF-β1 (rTGF-β1; 1 ng/ml) for 30 min. Cells were washed, treated with SB431542 (2 µM), and harvest at each time point to compared pSmad2 disappearance speed by WB. **(E)** Real-time PCR was performed using the same cDNA as in Figure 3F. **(F)** Mink lung epithelial cells (MLECs) stably harboring the TGF-β-responsive firefly luciferase (fLuc) construct were transfected with a Renilla luciferase (rLuc) construct and then seeded onto GFP- or DLX5-HMFs that had been treated with DOX (0.5 µg/ml) for three days. Luciferase activity was measured after 18 h of co-culture. fLuc/rLuc values relative to the GFP control are shown. *p*-values were determined by Tukey’s test (n = 6 to 9 from 3 independent experiments). **(G)** Immunoprecipitation (IP)-WB for GFP- or DLX5-HMFs. All cells were treated with DOX (0.5 µg/ml) for 48 h. rTGF-β1 treatment (1 ng/ml) was for 1 h prior to cell harvest. DNase treatment for IP samples was performed before the washing step. **(H)** Real-time PCR analysis of GFP- or DLX5-HMFs treated with DOX for 48 h and subsequently stimulated with rTGF-β1 (1 ng/ml) and DOX for 24 h. *p*-values were determined by Tukey’s test (n = 3). **(I)** TGF-β activity measured using the TGF-β reporter MLECs. *p*-value was determined by Mann–Whitney U test (n = 3 (fresh medium) and 4 (C.M.)). **(J)** Real-time PCR for GFP- or DLX5-HMFs which were treated with indicated media containing DOX (0.5 µg/ml) for 4 days. *p*-values were determined by Tukey’s test (n = 4). **(K)** HMFs were treated with rTGF-β1 (1 ng/ml) for 3 h prior to real-time PCR. *p*-value was determined by Welch’s t-test (n = 4). **(L)** Real-time PCR was performed using the same cDNA as in (C). The primers were designed in the 5’ or 3’ untranslated regions of *DLX5* to avoid amplifying exogenous *FLAG-DLX5* cDNA, which lacks these regions. *p*-values were determined by Tukey’s test (n = 4). **(M)** DLX5 promotes LRRC15 and POSTN expression by upregulating canonical TGF-β signaling and forming a complex with Smad2/3/4. Exogenous TGF-β induces DLX5, which in turn promotes its own expression in HMFs, possibly establishing a positive feedback loop. TGF-β activation downstream of DLX5 presumably promotes this loop. Error bars, SE; \**p* < 0.05, \*\**p* < 0.01 and \*\*\**p* < 0.001 (C, F, H-L).

Given the elevated canonical TGF-β signaling in DLX5-HMFs, we determined whether DLX5 increases the stability of pSmad2 protein. To address this issue, we evaluated the disappearance of pSmad2 protein following SB431542 treatment in DLX5-HMFs, which had been treated with recombinant TGF-β1 (rTGF-β1). pSmad2 levels were indeed comparable between DLX5-HMFs and GFP-HMFs for 2 h after the SB431542 treatment (Fig. 4D), suggesting that pSmad2 stability is not influenced by DLX5 in DLX5-HMFs.

We next reasoned that DLX5 may promote pSmad2 activation. We therefore checked TGF-β expression levels in DLX5-HMFs. *TGFB1*, *TGFB2* and *TGFB3* mRNA expression levels tended to be upregulated compared to those in GFP-HMFs (Fig. 4E). The TGF-β activity in media conditioned by DLX5- and GFP-HMFs was undetectable when measured by a TGF-β-responsive reporter system (Fig. EV4C). In contrast, significantly higher TGF-β activity was detected when reporter cells were directly co-cultured with DLX5-HMFs than with GFP-HMFs (Fig. 4F). This increase was not attributable to differences in the number of HMFs grown on culture dishes (Fig. EV4D) or reporter cells attached to fibroblasts (see the Figure legend and Methods), suggesting direct interaction with reporter cells to be required for the detection of increased TGF-β activity in DLX5-HMFs. Taken together, these findings suggest that active TGF-β produced by DLX5-HMFs contributes, at least in part, to pSmad2 activation in these fibroblasts.

We also reasoned that DLX5 may interact with an R-Smad complex. To address this speculation, we performed an immunoprecipitation assay using an antibody against FLAG tag fused to DLX5 in DLX5-HMFs. As anticipated, Smad2, Smad3 and Smad4 proteins were immunoprecipitated with the DLX5 protein (Fig. 4G). Consistently, the interaction of DLX5 with Smad2 and Smad4 has been reported in a transient co-transfection system (Maira *et al*, 2010). DNase treatment of the precipitated samples hardly interfered with the interaction between DLX5 and R-Smad proteins, suggesting their interaction to be regulated in a DNA-independent manner (Fig. 4G). The interaction with Smad2/3/4 was further increased by treatment with rTGF-β1 for 1 h prior to immunoprecipitation (Fig. 4G). Treatment with rTGF-β1 (Fig. 4H) or medium conditioned by active TGF-β-producing MCF7-*ras* breast cancer cells (Fig. 4I) also enhanced the increased *POSTN* and *LRRC15* mRNA expression levels in DLX5-HMFs (Fig. 4J). Taken together, these data suggest that DLX5 interacts with an R-Smad complex to induce TGF-β signaling and myCAF state in collaboration with active TGF-β in HMFs.

### TGF-β1 initiates DLX5 expression, establishing a positive feedback loop that sustains DLX5 expression in fibroblasts

Given the increased expression of DLX5 in myCAFs, we investigated how DLX5 expression is induced and maintained in these cells. HMFs were thus treated with rTGF-β1, followed by measurement of DLX5 expression. *DLX5* mRNA expression was induced in these cells by 22.3-fold, compared with untreated cells (Fig. 4K).

We also reasoned that DLX5 expression may be induced by DLX5 via autoregulation, because an enhancer region within the *DLX5/6* locus is reportedly bound by DLX5 itself (Maira *et al*., 2010; Zerucha *et al*, 2000). To address this speculation, endogenous *DLX5* mRNA expression was measured by real-time PCR analysis using primers specifically designed on *DLX5*-5’ and -3’ untranslated regions. Expression levels of the endogenous *DLX5* mRNAs were upregulated by 28.3- and 42.1-fold, respectively, in DLX5-HMFs relative to GFP-HMFs (Fig. 4L). SB431542 treatment significantly attenuated the *DLX5* induction, suggesting that the positive autoregulatory feedback loop of DLX5 is mediated by TGF-β signaling. Taken together with our earlier findings, these results demonstrate that DLX5, initially induced by TGF-β1, not only forms a complex with R-Smads to enhance canonical TGF-β signaling and promote the myCAF state, but also upregulates its own expression through a positive feedback loop in HMFs (Fig. 4M).

### DLX5 globally induces the myCAF state via binding to regulatory regions in target genes

To further elucidate molecular insights underlying the DLX5-primed myCAF state, we examined DLX5-regulated genes using genome-wide and bulk RNA-seq data of DLX5-HMFs. Gene set enrichment analysis (GSEA) confirmed DLX5-induced gene programs to be associated with ossification and the TGF-β signaling pathway (Fig. 5A). Notably, there was a significant enrichment of the CAF2 cluster but not the MF2/CAF1 cluster (Fig. 5B), indicating that DLX5 mediates myCAF-like gene expression programs in human breast cancers. Gene sets related to ECM were also enriched in DLX5-HMFs (Fig. 5C). SB431542 treatment significantly attenuated expression of the ECM genes, suggesting their TGF-β signaling-dependent regulation (Fig. 5D).

**Figure 5.**
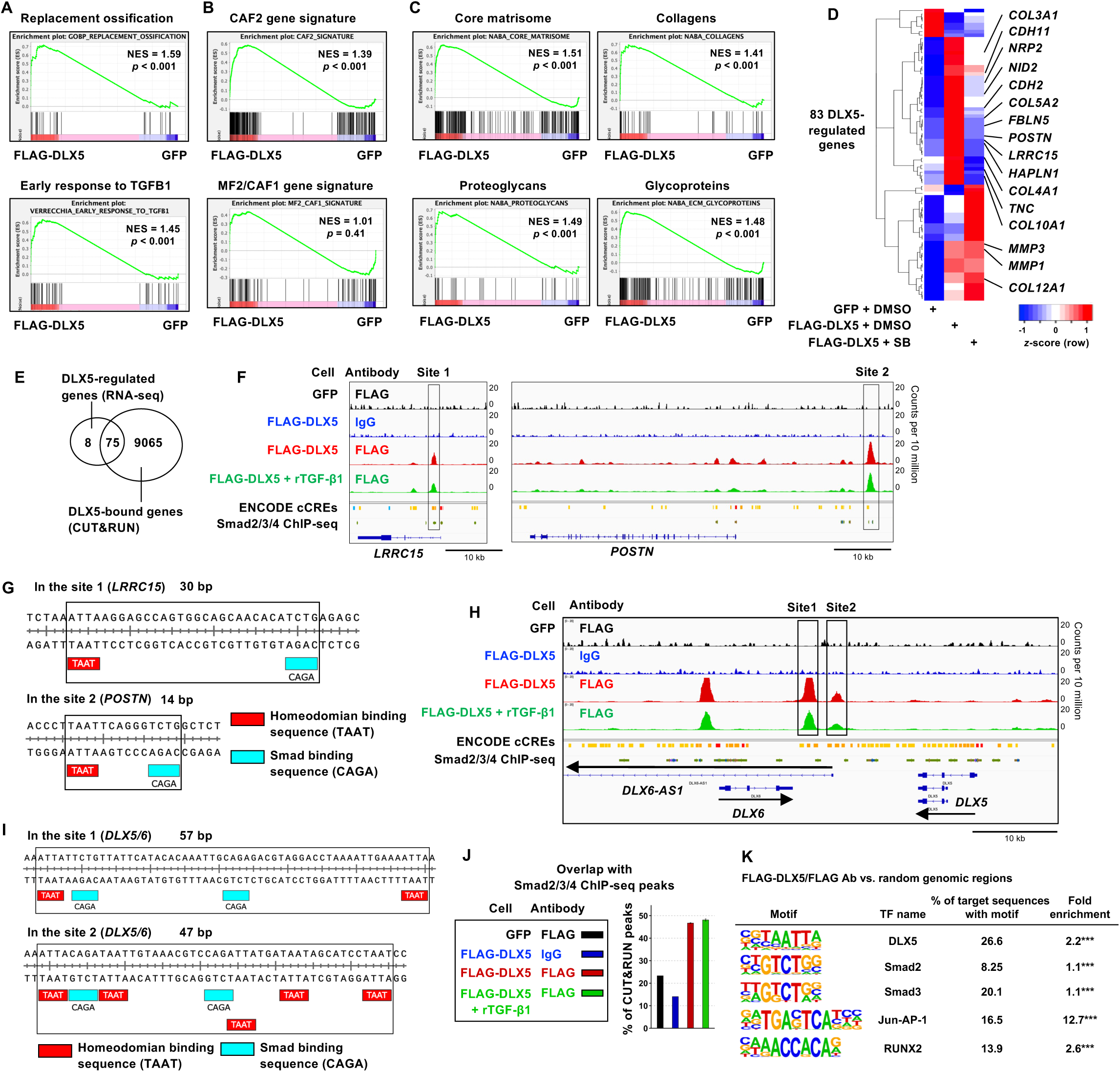
Cooperative occupancy of myCAF marker genes and the *DLX5* locus by the DLX5-R-Smad complex. **(A-C)** A subset of the total RNA harvested for Figure 4C was subjected to RNA-seq analysis (n = 3 per condition). Gene set enrichment analysis (GSEA) for DMSO-treated samples. NES, normalized enrichment score. *p*-values were determined by the permutation test. **(D)** Differentially expressed genes between DMSO-treated DLX5-HMFs and GFP-HMFs were identified (genes with an adjusted *p*-value < 0.05 and Transcripts Per Million > 10; Supplementary Table 3) and shown as *z*-scores. **(E)** GFP- or DLX5-HMFs were treated with DOX (0.5 µg/ml) for 3 days and subjected to CUT&RUN with an anti-FLAG antibody. rTGF-β1 (1 ng/ml) was added 24 h prior to CUT&RUN. Venn diagram showing the overlap between DLX5-bound genes and DLX5-regulated genes identified in (D). **(F)** Genome browser images. Orange to yellow, red, and blue bars on the candidate cis-regulatory element (cCRE) track indicate enhancer-like, promoter-like, and CTCF-bound regions, respectively. Smad2/3/4 chromatin immunoprecipitation sequencing (ChIP-seq) data were obtained from ReMap2022. **(G)** Representative genomic sequences containing TAAT and CAGA motifs in close proximity. **(H, I)** The *DLX5/6* locus was shown as in (F) and (G). **(J)** Proportions of CUT&RUN peaks obtained with FLAG antibody or IgG that co-localized with Smad2/3/4 ChIP-seq peaks. GFP/FLAG Ab (n = 1), FLAG-DLX5/IgG (n = 1), FLAG-DLX5/FLAG Ab (n = 2) and FLAG-DLX5 + rTGF-β1/FLAG Ab (n = 2). Error bars, S.D. **(K)** Motif enrichment analysis for FLAG-DLX5/FLAG Ab sample. *p*-values were calculated using the hypergeometric test and adjusted for multiple testing using Benjamini–Hochberg method in HOMER (\*\*\**q* < 0.001).

To investigate whether DLX5 directly regulates myCAF markers and ECM genes, we performed CUT&RUN for DLX5-HMFs treated with or without rTGF-β1. We identified 75 DLX5 direct target genes including *LRRC15* and *POSTN* among 83 differentially expressed genes extracted from bulk RNA-seq data (Fig. 5E). DLX5 CUT&RUN peaks overlapped with publicly available Smad2/3/4 chromatin immunoprecipitation sequencing (ChIP-seq) peaks and with enhancer regions predicted from active histone modifications in the ENCODE project around *LRRC15* and *POSTN* loci (Fig. 5F). Core binding sequences for DLX5 (TAAT) and Smad (CAGA) were also located closely in the representative target genes (Fig. 5G), suggesting cooperative genomic occupancy between DLX5 and R-Smads.

DLX5 binding peaks were also detected at the *DLX5/6* locus, which contains TAAT and CAGA sites within each peak (Fig. 5H, I). These sites 1 and 2 include the evolutionarily conserved enhancer regions bound and activated by DLX5, known as hI56i and hI56ii, respectively (Maira *et al*., 2010; Zerucha *et al*., 2000), suggesting the autoregulation of DLX5, consistent with earlier data (Fig. 4L, 5H, I). We further revealed that almost 47% of regions bound by DLX5 overlapped with Smad2/3/4 ChIP-seq peaks through the genome (Fig. 5J). However, motif enrichment analysis showed that DLX5, Jun-AP-1 and RUNX2 motifs were enriched in DLX5-bound regions, whereas Smad2/3 motifs were not consistently enriched across different analysis conditions (0.5-1.1-fold) (Fig. 5K, EV4E). The discrepancy between these results may reflect differences in the underlying approaches: the former relies on Smad2/3/4 ChIP-seq data, which may include a substantial number of indirect binding sites lacking the consensus Smad-binding motifs, whereas the latter identifies potential direct DNA-binding sites based on motif sequences. Taken together, these findings suggest that the DLX5-R-Smad complex co-occupies at least the *POSTN*, *LRRC15* and *DLX5* loci, thereby promoting the myCAF state and DLX5 expression.

### DLX5-HMFs promote the collective invasion of human breast cancer cells

Given that TGF-β signaling and the myCAF state were increased in DLX5-HMFs, we sought to determine whether these fibroblasts impact tumor progression. To address this issue, we performed a 3D co-culture assay employing DLX5-HMFs and MCF10A-DCIS.com breast cancer cells, modified to express tdTomato, blasticidin resistance and firefly luciferase (DCIStb cells) (Hu *et al*, 2008; Matsumura *et al*., 2019) (Fig. 6A). We observed significantly increased invasive protrusions formed by DCIStb cells (indicated by arrowheads) under the DOX(+) condition for 4 days compared to the DOX(-) condition (Fig. 6B).

**Figure 6.**
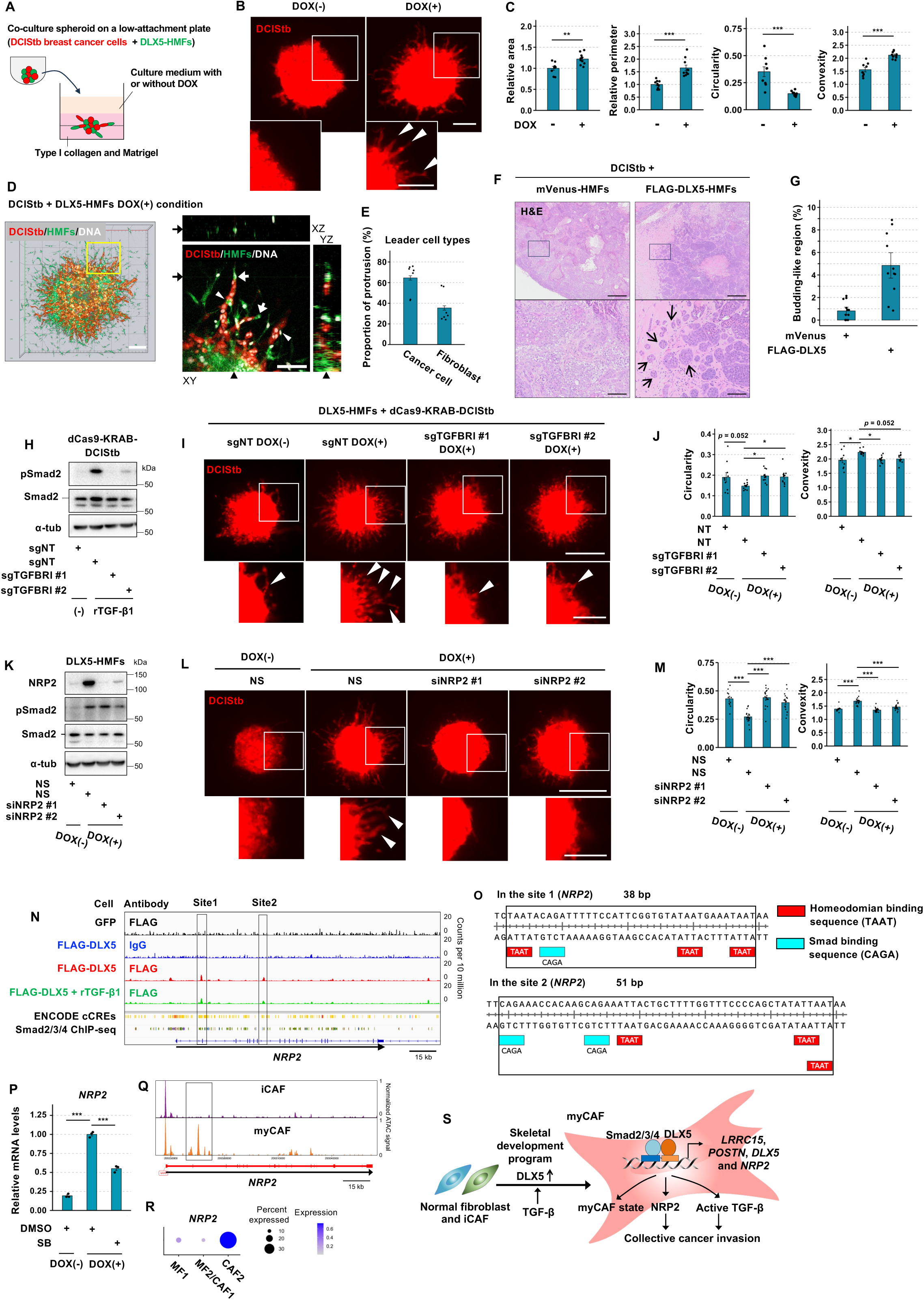
DLX5-HMFs promote breast cancer invasion via active TGF-β production and NRP2 induction. **(A)** Schematic representation of the co-culture spheroid assay. **(B)** Images obtained by epifluorescence microscopy at 4 days from gel embedding. Scale bars, 250 µm. Arrowheads in (B), (I), and (L) indicate cancer cell invasion. **(C)** Morphometric parameters for tdTomato. See Fig. EV5A. *p*-values were determined by Student’s t-test (n = 9 and 10 from three independent experiments). **(D)** Left, 3D-rendered image obtained by confocal microscopy. Scale bar, 200 µm. Right, enlarged optical sections of the inset region. Fibroblasts as leader cells (arrows) and follower cells (arrowheads). Scale bar, 100 µm. **(E)** Categorization of leader and follower cells. Seven spheroids (DOX(+)) from three independent experiments. Dots, spheroids. 15-32 protrusions per each spheroid were evaluated. Error bars, SE. **(F)** H&E images for sections prepared from xenograft tumors at day 29. Scale bars, 500 µm (top panel) and 250 µm (bottom panel). Arrows, invading cancer cells forming budding-like structures. **(G)** Budding-like regions was quantified. *p*-value was determined by Mann–Whitney U test (n = 12 (mVenus) and 11 (DLX5)). **(H)** DCIStb cells expressing both dCas9-KRAB and indicated sgRNA were treated with rTGF-β1 for 30 min, followed by WB. NT, non-targeting sgRNA. **(I, J)** These cells were subjected to the 3D co-culture assay with FLAG-DLX5-HMFs with or without DOX (0.5 µg/ml). Images and morphometric parameters for tdTomato at day 4. Scale bars, 500 µm and 250 µm (inset). *p*-values were determined by Dunnett’s t-test (n = 10 to 12 from three independent experiments). **(K)** DLX5-HMFs were transfected with siRNA and treated with DOX (0.5 µg/ml) for 72 h starting the following day, followed by WB. NS, non-silencing siRNA. **(L, M)** Co-culture spheroid assay using DCIStb cells and DLX5-HMFs transfected with the indicated siRNA 2 days prior to the assay. DOX treatment was started on the day of gel embedding (0.5 µg/ml). Images and morphometric parameters for tdTomato at day 3. Scale bars, 500 µm and 250 µm (inset). *p*-values were determined by Dunnett’s t-test (n = 12 to 18 from three independent experiments). **(N, O)** CUT&RUN data, same as in Figure 5F and G. **(P)** Real-time PCR analysis of FLAG-DLX5-HMFs treated with the indicated chemicals for 48 h. SB431542 (2 μM) and DOX (0.5 μg/ml). *p*-values were determined by Dunnett’s t-test (n = 3). **(Q)** Pseudobulk clusters were generated using cells of cluster 5 (myCAFs) and cluster 4 and 6 (iCAFs) in Figure 2G. The box highlights a region with increased ATAC signal in myCAFs. **(R)** *NRP2* mRNA expression in the scRNA-seq dataset shown in Figure 1E. **(S)** myCAFs emerge from normal mammary fibroblasts and iCAFs. DLX5, a key regulator of the skeletal developmental program, is activated in myCAFs at both epigenetic and transcriptional levels. DLX5 expression is initiated by TGF-β1 in fibroblasts, enabling DLX5 to form a complex with Smad2/3/4 and cooperatively promote canonical TGF-β signaling, thereby inducing the myCAF state, active TGF-β production and NRP2 expression. DLX5-primed myCAFs also promote breast cancer invasion through paracrine TGF-β signaling and NRP2 expression. Error bars, SE; *p < 0.05, **p < 0.01 and ***p < 0.001 (C, E, G, J, M, P).

To evaluate more precisely the protrusion-forming collective invasion, we employed various metrics including area, perimeter, circularity (4π×spheroid area/(spheroid perimeter)^2^) and convexity (spheroid perimeter/convex hull perimeter) of tdTomato fluorescence (Fig. EV5A). All these four parameters showed significantly increased tumor protrusions in the DOX(+) condition (Fig. 6C). DLX5-HMFs also led tumor cells at 35% of tumor protrusions under confocal microscopy in the DOX(+) condition (Fig. 6D, E), consistent with a previous report that demonstrated the fibroblast-driven tumor protrusion (Gaggioli *et al*, 2007).

To further examine roles of DLX5-HMFs in tumors, we coinjected HMFs constitutively expressing FLAG-DLX5 (pCDH-DLX5-HMFs) with DCIStb cells subcutaneously into immunodeficient mice. Expression levels of pSmad2, LRRC15 and POSTN proteins were substantially upregulated in these fibroblasts prior to injection (Fig. EV5B). Of note, we observed a 6.0-fold increase in tumor budding-like structures, indicative to invading tumor cell clusters in tumors containing pCDH-DLX5-HMFs relative to the control pCDH-mVenus-HMFs (Fig. 6F, G). Primary tumor weight was comparable between these two groups (Fig. EV5C). Collectively, these findings demonstrate that DLX5-HMFs can promote tumor protrusions in human breast tumors.

### Paracrine TGF-β signaling mediates collective cancer invasion driven by DLX5-HMFs

Given the pro-invasive phenotype and active TGF-β production (Fig. 4F) in DLX5-HMFs, we speculated that paracrine TGF-β signaling from DLX5-HMFs may influence invasion of apposed carcinoma cells. To assess this speculation, we generated two different single-guide RNAs (sgRNAs) suitable for CRISPR interference targeting *TGFBRI*, which inhibited *TGFBRI* mRNA expression in DCIStb cells by 90.3-99.3%, respectively, compared to the effect of the control non-targeting (NT) sgRNA (Fig. EV5D). *TGFBRI* inhibition by sgRNAs also attenuated pSmad2 levels in DCIStb cells treated with rTGF-β1 by 62.6-85.1%, indicating reduced responsive to external TGF-β stimulation (Fig. 6H). When TGFBRI-sgRNA-expressing DCIStb cells were co-cultured with DLX5-HMFs under the DOX(+) condition, we found significantly suppressed tumor protrusions, based on evaluation using the aforementioned parameters (Fig. 6I, J, EV5E). These findings indicate that paracrine TGF-β signaling from DLX5-HMFs contributes to promotion of tumor protrusions.

### Neuropilin-2 (NRP2), a target gene of DLX5, mediates tumor invasion driven by myCAFs

Since inhibition of paracrine stromal TGF-β signaling suppressed tumor protrusions but the effect was not strong, we sought to identify DLX5 target genes that mediate the pro-invasive program. As *NRP2* is included in the TGF-β-dependent DLX5-regulated genes and in the cardiovascular signature of the CAF2 cluster, as mentioned earlier (Fig. 2D, 5D), we decided to investigate the role of NRP2. NRP2 is a co-receptor for various signal receptors including a TGF-β receptor, and its expression is also positively regulated by TGF-β signaling (Islam *et al*, 2022; Wittmann *et al*, 2015). However, roles of NRP2 on myCAFs are poorly understood. Thus, we generated two different siRNAs against *NRP2*, which suppressed NRP2 protein by 92.8-95.2% and *NRP2* mRNA expression by 84.1-89.1%, respectively, compared to the effect of the control non-silencing (NS) siRNA in DLX5-HMFs (Fig. 6K, EV5F). We transfected these siRNAs before the 3D co-culture assay using DCIStb cells. Of note, inhibition of NRP2 expression strongly attenuated tumor protrusion (Fig. 6L, M, EV5G), indicating that NRP2 is required for DLX5-HMFs to promote tumor protrusion.

We also investigated regulatory mechanisms of NRP2 induction by DLX5 using CUT&RUN data for DLX5-HMFs. We found that DLX5 binds to the *NRP2* locus, which contains putative Smad binding sites (Fig. 6N, O). The DLX5-induced upregulation of *NRP2* mRNA was also significantly attenuated by SB431542 treatment (Fig. 5D, 6P), indicating DLX5-induced NRP2 upregulation is regulated by TGF-β signaling. Conversely, NRP2 inhibition by siRNA did not influence the pSmad2 level in DLX5-HMFs (Fig. 6K). Taken together, these observations suggest that NRP2 expression is induced by DLX5 via TGF-β-Smad2/3 signaling, but NRP2 hardly affects the canonical TGF-β-Smad2/3 signaling in these fibroblasts.

We also investigated stromal NRP2 expression in breast cancer patients. Chromatin accessibility at the *NRP2* locus and *NRP2* mRNA expression were increased in myCAFs but not iCAFs of breast cancer patients, as gauged by scATAC-seq and scRNA-seq analyses, respectively (Fig. 6Q, R, EV5H), indicating NRP2 to be enriched in human breast myCAFs. Taken together, these data suggest that myCAFs promote tumor protrusion presumably through NRP2 induced by DLX5 in collaboration with canonical TGF-β signaling in breast cancer patients (Fig. 6S).

## Discussion

### DLX5 induces tumor-promoting myCAFs in collaboration with TGF-β signaling

myCAFs, a major subtype of CAFs, are predominantly present in human breast cancers to influence hallmarks of cancer, including tumor invasion (Bartoschek *et al*., 2018; Croizer *et al*., 2024; Liu *et al*., 2026; Mezawa & Orimo, 2022). myCAFs are induced by activation of TGF-β signaling during tumor progression. However, molecular mechanisms by which TGF-β signaling impacts myCAF-specific transcriptional programs remain poorly understood. In this study, using a relatively large integrated scRNA-seq dataset of human breast tissues, we demonstrate that myCAFs activate skeletal and cardiovascular developmental programs that are associated with poorer prognosis in breast cancer patients.

DLX5, a central TF for the skeletal developmental program, is also enriched in myCAFs. DLX5 activates TGF-β-Smad2/3 signaling to induce the myCAF state in human mammary fibroblasts. DLX5-primed myCAFs have the ability to promote the collective invasion of human breast cancer cells. Mechanistically, DLX5 expression is initiated by TGF-β1 treatment in human mammary fibroblasts, and DLX5 interacts with a Smad2/3/4 complex, enabling cooperative genomic occupancy of DLX5 and R-Smads to promote the myCAF state and active TGF-β production (Fig. 6S). DLX5-primed myCAFs also induce paracrine TGF-β signaling and NRP2 expression to promote collective breast cancer invasion (Fig. 6S). Taken together, the present study highlights an unappreciated role of DLX5 to give rise to myCAFs by activating canonical TGF-β signaling, thereby contributing to human breast cancer invasion.

### Skeletal and cardiovascular development programs activated in human breast myCAFs

The skeletal developmental program in the CAF2 cluster, in which myCAFs are highly enriched, involves genes related to ECM components and ECM-remodeling enzymes. These include *COL11A1*, *COL10A1*, *BMP1* and *POSTN*, which are essential for development or homeostasis of the skeletal system (Muir *et al*, 2014; Richards *et al*, 1996; Shen, 2005; Wishman *et al*, 2025). LRRC15 is also expressed in osteoblasts (Purcell *et al*, 2018). On the other hand, we noted lower expression levels of *COL2A1*, alkaline phosphatase (*ALPL*) and integrin binding sialoprotein (*IBSP*) (Fig. EV4F), which are markers for mature chondrocytes and osteoblasts (Amarasekara *et al*, 2021). These findings suggest that myCAFs might partially transdifferentiate into osteoblastic and chondrocytic lineages to contribute to ECM production and remodeling. Given that perivascular cells, osteoblasts, chondrocytes and fibroblasts are developmentally related mesenchymal cells, ectopic activation of skeletal and cardiovascular developmental genes, including DLX5 and RUNX2, in myCAFs could be considered as a type of lineage perturbation, similar to unlocking lineage plasticity in cancer cells (Lendahl *et al*, 2022; Tirosh & Suva, 2024).

### Cooperative genomic occupancy by the DLX5-Smad2/3/4 complex in myCAFs

DLX5 interacts with Smad2/3/4 proteins and the resulting complex shares binding sites for DLX5 and R-Smads to cooperatively induce myCAF marker genes. rTGF-β1 treatment increased the abundance of the DLX5-Smad2/3/4 protein complex (Fig. 4G) and further enhanced *LRRC15* and *POSTN* mRNA expression in DLX5-HMFs (Fig. 4H). However, the CUT&RUN peak intensities did not increase accordingly (Fig. 5F). This may be due to a sufficient level of the DLX5-R-Smad complex to occupy its binding sites without supplementation with rTGF-β1, since the active TGF-β level is increased in DLX5-HMFs (Fig. 4F). The reason that rTGF-β1 treatment further enhances *LRRC15* and *POSTN* mRNA expression in DLX5-HMFs without increasing DLX5 occupancy at these loci remains unclear. However, we speculate that rTGF-β1 treatment increases the DLX5-R-Smad complex, thereby possibly promoting the formation of transactivation hubs through interactions between the low-complexity domains of Smad4 and DLX5 (Chong *et al*, 2018; Li *et al*, 2024). Our structural analyses using AlphaFold3 and PrDOS also predicted that DLX5 contains intrinsically disordered regions (IDRs) that often provide the molecular basis for the formation of transactivation hubs (Fig. EV5I, J). Determining whether transactivation hubs are formed through interactions between these IDRs to enhance transcriptional activity would be an intriguing direction for future investigation.

Motif enrichment analysis showed substantial enrichment of Jun-AP-1 and RUNX2 motifs in DLX5-bound genomic regions (Fig. 5K, EV4E). *RUNX2* mRNA expression and RUNX2 activity were also more increased in the CAF2 cluster compared with the MF1 and MF2/CAF1 clusters (Fig. 3B, C). Therefore, these TFs may also contribute to myCAF formation, potentially through interactions with the DLX5-R-Smad complex. Previous reports describing interactions between RUNX2 and DLX5 and between AP-1 and SMAD3/4 (Liberati et al., 1999; Roca et al., 2005) support this possibility.

### DLX5-primed myCAFs promote collective invasion of breast cancer cells

DLX5-HMFs promote tumor cell invasion by leading human breast cancer cell clusters in 3D culture system (Fig. 6A-D). TGF-β activity (Fig. 4F) and NRP2 expression (Fig. 6K, P) are increased in DLX5-HMFs. TGFBRI inhibition in cancer cells and NRP2 suppression in DLX5-HMFs indeed significantly attenuate collective breast cancer cell invasion in the 3D culture system (Fig. 6H-M). These observations therefore indicate that increased paracrine TGF-β signaling and NRP2 expression in DLX5-HMFs mediate tumor cell invasion.

Meanwhile, we observe a higher level of GFP fluorescence in tumor spheroids admixed with GFP-expressing DLX5-HMFs treated with DOX relative to no DOX treatment (Fig. EV5K), indicating the greater number of fibroblasts present in the spheroids with DLX5-HMFs. As NRP2 is a coreceptor for signaling pathways of various growth factors (Islam *et al*., 2022), we speculated that NRP2 may promote the survival and/or growth of fibroblasts present in spheroids. Contrary to our expectations, inhibition of NRP2 expression by two different siRNAs failed to consistently decrease the number of DLX5-HMFs (Fig. EV5L). These findings thus suggest that the pro-invasive phenotype by stromal NRP2 is independent of the control of fibroblast survival and/or growth. Cancer cell-derived signals, such as VEGFs and semaphorins, may act on NRP2 expressed on DLX5-HMFs, allowing these fibroblasts to produce as-yet-unidentified invasion-promoting factors.

Collectively, the present study highlights novel roles for DLX5 in orchestrating a skeletal developmental program that promotes myCAF formation through cooperation with canonical TGF-β signaling. DLX5-primed myCAFs, in turn, promote breast tumor invasion via paracrine TGF-β signaling and NRP2 induction.

## Methods

### Preparation and analyses of integrated scRNA-seq data

Publicly available scRNA-seq data for human mammary tissue and breast cancer were obtained from NCBI GEO and EMBL-EBI BioStudies: GSE198732 (Murrow *et al*., 2022), GSE180878 (Gray *et al*., 2022), GSE164898 (Bhat-Nakshatri *et al*., 2021), GSE161529 (Pal *et al*., 2021), E-MTAB-10607 (Tietscher *et al*., 2023), GSE176078 (Wu *et al*., 2021), GSE199515 (Delgado *et al*., 2024). Full patient list and clinical background were described in Supplementary Table 1. Data from patients who had undergone prophylactic surgery were omitted if the information was provided in the original papers. Data processing and analyses were performed in Seurat (ver. 5) (Hao *et al*, 2024) and R (ver. 4.3.2). Cells in each dataset of 132 donors were filtered with number of detected genes, RNA and mitochondria derived RNA to remove poor quality cells, and were then subjected to DoubletFinder (McGinnis *et al*, 2019) for depleting expected doublets. Data of all patients were merged and processed with NormalizeData, FindVariableFeatures and RunPCA with default settings. Next, Harmony (ver. 1.2.0) (Korsunsky *et al*, 2019) and JoinLayers were employed to correct batch effects and integration, respectively. The integrated data were subjected to FindNeighbors, FindClusters and RunUMAP with default settings for clustering and dimensional reduction. For assignment of cell types, *EPCAM*, *PTPRC*, *PECAM1* and *MCAM* were used for epithelial cells, immune cells, ECs and PVs, respectively. *PDGFRA*, *DCN* and *COL1A1* were used for fibroblast markers.

After subclustering of fibroblasts and PVs, we noted that some small clusters (clusters 17, 18 and 22 in Figure EV1F) contained cells expressing keratin genes and either *COL1A1* or *PTPRC.* We removed them from analyses due to the possible doublets. Clusters which did not express any major lineage marker except for *VIM* (clusters 16 and 19) and cells having read counts for any of *KRT8*, *KRT14*, *KRT17* or *KRT18* were also removed for accurately focusing on fibroblasts and PVs in this study. As a consequence, we obtained 39,058 cells consisting of 18 clusters (Fig. 1C-E). The procedure of grouping cells into cluster types (MF1, 2, CAF1, 2, MP, SMC and CAP) was written in the results section. Normalized read counts were presented by heatmaps on balloon plots and dimensional reduction plots. The percentage of expressed cells was calculated by tallying cells with any level of read count for a gene of interest. For the pseudotime trajectory analysis, Monocle3 was used with default settings (Cao *et al*, 2019). Although Monocle3 identified three partitions, we focused on two of them: the two partitions enriched for fibroblasts or PVs because the other partition was tiny.

To obtain gene signatures of the MF2/CAF1 clusters and CAF2 clusters, FindMarkers was used with default setting but configured to identify only positive markers expressed in more than 25% of cells in the analyzed cluster. The genes having a p < 0.05 and a fold change in an expression rate > 2 (CAF2 over MF2/CAF1 and MF1, or MF2/CAF1 over CAF2 and MF1) were included in the gene signature. They were subjected to GO analysis in PANTHER.db (ver. 18.0). Enrichment analysis for human diseases and mouse phenotypes was performed with PhenoExam (Cisterna *et al*, 2022).

To infer activity of TFs or signal pathways, decoupleR (ver. 2.9.7) (Badia et al, 2022) was applied to randomly subsampled sets of 4,000 and 8,000 cells, respectively. The 20 most variable TFs were selected according to standard deviations of each TF. The scRNA-seq data of breast CAF-S1 was kindly gifted from Dr. Mechta-Grigoriou (Institut Curie, Paris) (Kieffer *et al*., 2020). Expression levels and TF activity were analyzed with Seurat and decoupleR, respectively.

### Analysis of TCGA breast cancer data

Survival and expression data for TCGA Breast cancers were obtained from UCSC Xena. To evaluate the impact of CAFs on patients’ survival time, we selected genes whose expression rates in fibroblasts were more than twofold higher than those in each of the other cell types shown in Fig. 1B, using data from the cancer condition. Expression levels of the CAF specific cardiovascular and skeletal developmental genes (highlighted by a gray background in Fig. 2D) were transformed into a single score using GSVA (ver. 1.50.0). Patients were separated into two groups with the median, and then subjected to Kaplan-Meier plot analysis and the Log-Rank test in R package Survival (ver. 3.5.7).

### Data analyses of scATAC-seq of human breast cancers

The scATAC-seq data of 16 breast cancer patients published by us were reanalyzed with ArchR (Granja *et al*., 2021; Kumegawa *et al*., 2022). These included 11, 2 and 3 breast cancer cases of the luminal, human epidermal growth factor receptor 2 (HER2)-positive, and triple-negative subtypes, respectively. CAFs and perivascular cells (PVs) were identified from all cells based on gene scores for *COL1A1* and *PDGFRA* (CAFs) and MCAM (PVs), followed by subclustering. To distinguish CAFs from PVs, the following marker genes were used: *COL1A1*, *PDGFRA*, *DCN*, and *LUM* for fibroblasts, and *MCAM*, *RGS5*, *CSPG4*, and *MYH11* for PVs. Gene activity score, module score and pseudobulk analyses were performed in ArchR. Module scoring and pseudobulk analyses were performed using ArchR’s addModuleScore and plotBrowserTrack functions.

### Cell culture

A human mammary fibroblast cell line (218TGpp: HMFs) was established in our previous study (Kojima *et al*., 2010) and maintained in DMEM high glucose supplemented with 10% of fetal bovine serum (FBS) and 1% of penicillin-streptomycin mixture. MCF10ADCIS.com cells that were transduced with pWZL-tdTomato-blast in our previous study (Matsumura *et al*., 2019) were maintained in DMEM/F12 supplemented with 5% FBS and 1% penicillin–streptomycin. These cells were also transduced with pCDH-EF1-Luc2-IRES-puro. All cell lines used in the present study were confirmed not to be infected with mycoplasma. For siRNA transfection experiments, 1 × 10 fibroblasts were seeded in 6-well plates, followed by siRNA transfection the next day. Eight picomoles of siRNA and 2 µl of Lipofectamine™ RNAiMAX Transfection Reagent (Thermo Fisher Scientific, 13778150) were mixed in 200 µl of Opti-MEM (Thermo Fisher Scientific, 31985062). The mixture was incubated for 20 min and then applied to the fibroblasts for 6 h.

### Bulk RNA-seq

218TGpp-CSIV-GFP and -FLAG-DLX5 were treated with DOX (0.5 µg/ml) and SB431542 (2 µM) or DMSO (1:5000) for 4 days. Total RNA was prepared with NucleoSpin RNA (Takara). Library preparation and sequencing were outsourced to KOTAI Biotechnologies, Inc. (Osaka, Japan). RNA integrity was evaluated using the Agilent fragment analyzer with the corresponding Kit (Agilent Technologies). RNA sequencing libraries were constructed using the NEBNext® Ultra™ II Directional RNA Library Prep Kit (Cat No. E7760, New England Biolabs) according to the manufacturer’s instructions. Briefly, polyadenylated mRNA was enriched from 400 ng of total RNA using oligo-dT magnetic beads. The mRNA was then fragmented and reverse-transcribed into first-strand cDNA using random hexamer primers. The second strand cDNA was synthesized using dUTP, instead of dTTP. The directional library was ready after end repair, A-tailing, adapter ligation, size selection, amplification, and purification. The quality and size distribution of the libraries were assessed using the AATI fragment analyzer (Agilent Technologies). The cyclization was performed using MGIEasy universal library conversion kit (App-A). Sequencing was performed on the DNBSEQ-T7 platform (MGI Tech.) using a paired-end 150 bp (PE150) sequencing strategy. The sequencing depth was targeted at 20 million reads per sample to ensure sufficient coverage for downstream analysis. FASTQ files were mapped to GRCh38, and Transcripts Per Million (TPM) and differential gene expression was obtained by RSEM and DESeq2. Genes meeting an adjusted *p* < 0.05 between 218TGpp-CSIV-GFP and -FLAG-DLX5, and TPM > 10 were defined as DLX5-regulated genes. GSEA (ver. 4.0.3) was performed on software obtained from Broad Institute with default settings.

### CUT&RUN

0.8×10^5^ fibroblasts were seeded onto 6 well plates and DOX (0.5 µg/ml) was supplemented at the following day. The culture medium was replaced with culture medium supplemented with 0.5% FCS and human rTGF-β1 (1 ng/ml) or vehicle (1 mg/ml BSA-1 mM HCl) on day 3 after cell seeding. After 24 h, CUT&RUN was performed using ChIC/CUT&RUN Assay Kit according to the manufacturer’s manual (Active Motif, 53180). Samples derived from 2×10^5^ fibroblasts were supplemented with 5,000 *Drosophila* nuclei (Active Motif, 53183) and incubated with 1 µg of anti-FLAG antibody (Sigma, clone M2) and 1 µg of spike-in antibody. Library preparation was performed with NEBNext Ultra II DNA Library Prep Kit for Illumina (NEB, E7645S) according to the manual. Next generation sequencing with Illumina NovaSeq X Plus was outsourced to Azenta, Inc. Those sequence reads were mapped onto GRCh38 and BDGP6 with Bowtie (ver. 2.4.1) and subjected to normalization by spike-in reads using samtools. Peak calling, Wiggle file generation, and motif enrichment analysis were performed using HOMER. Data from two replicates were averaged to generate Wiggle files and merged for motif analysis. Peaks were assigned to the single nearest gene (1000 kb maximal extension) using the GREAT (ver. 4.0.4). A series of Smad2/3/4 ChIP-seq datasets was obtained from ReMap2022 and merged into a single BED file to investigate overlap with FLAG-DLX5 CUT&RUN peaks using the mergePeaks command in HOMER. For motif enrichment analysis, the findMotifsGenome.pl command was run using HOMER.

### Plasmid vector construction and lentiviral infection

The cDNAs encoding FLAG-tagged human *DLX5* (kindly gifted from Dr. Joseph Testa at Fox Chase Cancer Center, Pennsylvania (Tan *et al*, 2008)), human *RUNX2* (Addgene, 143553) and *EGFP* were subcloned into pENTR1A (Addgene, 17398) followed by LR reaction to transfer to CSIV-TRE-Ubc-RfA-IRES-hKO1-T2A-rtTA (RIKEN BRC #RDB12878). hKO1 was replaced with TagBFP derived from pTagBFP-C vector (Evrogen) to generate CSIV-TRE-Ubc-RfA-IRES-TagBFP-T2A-rtTA. cDNA of *FLAG-DLX5* and *mVenus* were subcloned into pCDH-EF1-MCS-IRES-Neo vector (System Biosciences) using restriction enzymes. Preparation of lentivirus was performed as described previously with several modifications (Mezawa *et al*, 2023). Briefly, HEK293T cells were co-transfected with a lentiviral vector, pCMV-dR8.2dvpr and pCMV-VSVG using Fugene6 (Promega) or PEI MAX (Polysciences). The lentivirus was concentrated with ultracentrifugation and applied to 218TGpp cells. hKO1 or tagBFP expressing cells were sorted using MoFlo Astrios (BECKMAN COULTER) and expanded for experiments. For TGFBRI knockdown in DCIStb cells, the *GFP* gene in pLV hUbC-dCas9-KRAB-T2A-GFP (Addgene plasmid, 67620) was replaced with a hygromycin resistance gene using restriction enzyme cloning, followed by introducing a silent mutation that eliminated the BsmBI site within the hygromycin resistance gene. Oligonucleotides encoding non-targeting or *TGFBRI*-targeting sgRNAs were cloned into the BsmBI site downstream of the U6 promoter. Lentivirus was generated as mentioned above and used to transduce DCIStb cells, followed by hygromycin selection (50 µg/ml). The DNA sequences for sgRNA are listed in Supplementary Table 4.

### 3D co-culture assay

7,500 218TGpp cells transduced with CSIV-TRE-Ubc-FLAG-DLX5-IRES-tagBFP-T2A-rtTA and 2,500 FACS-sorted tdTomato-high DCIStb-Luc2 cells were seeded in a Nunclon™ Sphera™ 96-Well, Nunclon Sphera-Treated, U-Shaped-Bottom Microplate (Thermo Fisher Scientific, 174925) containing a 1:1 mixture of high-glucose DMEM and DMEM/F12 supplemented with 5% FBS and 0 or 0.5 µg/ml DOX. In the case of siRNA experiments, siRNA is transfected into those fibroblasts 48 h prior to seeding. After overnight incubation the spheroid was transferred onto 50 µl of a solidified ECM gel (3D Ready Atelocollagen DMEM-HG5 (KOKEN, 3D-HG05), Matrigel Growth Factor Reduced (Corning, 354230) and the same medium (but 0.5% FCS) were 12.5 µl, 6.25 µl and 31.25 µl, respectively) and incubated for 2.5 h to allow the spheroid to attach followed by mounting with 50 µl of the gel. After solidification, the same medium with or without DOX (0.7 µg/ml) was supplied. Fluorescence images were taken by Eclipse80i (Nikon) at day 3 or 4 from the gel embedding. Spheroid size and perimeter were obtained by using ImageJ. Next, spheroids were fixed with 4% PFA at room temperature for 1 h and washed with 1×PBS three times followed by incubation with SCALE VIEW S4 (Wako) containing TO-PRO-3 (1 µM) at 37 overnight. Z-stack images were prepared with LSM900 (Carl Zeiss). Preparation of 3D rendered or orthogonal images and the manual investigation of leader and follower cell types were performed in ZEN Pro (Carl Zeiss).

### Real-time PCR

RNA was prepared from cultured cells using NucleoSpin RNA (Takara). 500 ng of RNA was subjected to reverse transcription using PrimeScript RT reagent Kit (Takara) and used for real-time PCR using THUNDERBIRD Next SYBR qPCR Mix (TOYOBO) and QuantStudio3 (Thermo Fisher Scientific). Data were processed using the ΔΔCt method. The expression levels were normalized to those of glyceraldehyde 3-phosphate dehydrogenase (GAPDH). The primer sequences are presented in Supplementary Table 4.

### Western blotting (WB)

Cultured fibroblasts were lysed with 2×SDS sample buffer and protein concentration was determined by DC protein assay (Bio-Rad). SDS-PAGE and WB were performed using the same procedure as in our previous study (Mezawa *et al*, 2019).

### Immunoprecipitation-western blotting

1×10^6^ of 218TGpp CSIV-GFP or FLAG-DLX5 were seeded onto five 10 cm dishes. Those cells were treated with DOX (0.5 µg/ml) for 48 h prior to cell lysis with lysis buffer (25 mM Tris-HCl pH 7.5, 140 mM NaCl, 2.5 mM MgCl_2_, 1 mM EGTA, 0.5% Triton X-100 and 1×protease inhibitor cocktail). The cell lysate was incubated with 4 µg of anti-FLAG antibody (M2, Sigma) conjugated with 80 µg of protein G-FG beads (TAMAGAWA SEIKI) at 4 for 2 h. The beads were separated into two tubes and incubated with DNase reaction buffer with or without recombinant DNase (supplemented in NucleoSpin RNA kit, Takara) for 10 min at 37 to investigate a possibility of DNA-mediated co-precipitation. The beads were washed 4 times with wash buffer (cell lysis buffer which contained 0.1% Triton X-100 and 0.2×protease inhibitor) and proteins were eluted with 40 µl of 1.5×sample buffer supplemented with 2-mercaptoethanol at 95 for 3 min. 0.1% of input and 50% of IP samples were subjected to SDS-PAGE using an 8% polyacrylamide gel and WB. The antibodies used were written in Supplementary Table 4.

### Immunohistochemistry and double fluorescence immunostaining

Immunohistochemistry and double fluorescence immunostaining were performed as previously described with slight modifications (Mezawa *et al*., 2019). The use of formalin fixed paraffin embedded tissue specimens of breast cancers in this study was approved by the Juntendo University ethics review board. Sections were prepared from 7 cases positive for estrogen receptor (ER) and progesterone receptor (PR), and 2 cases negative for ER and PR but positive for HER2. Briefly, antigen retrieval was performed by using Decloaking Chamber™ NxGen in Tris-EDTA buffer at pH 9.0 for 40 minutes at 95°C. The slides were incubated with anti-DLX5 antibody at 4°C overnight. Secondary antibody (EnVision+ Single Reagents (HRP. Rabbit)) was incubated for 1 h at room temperature. Slides were incubated with 3,3′ diaminobenzidine solution (0.2 mg/ml) followed by hematoxylin staining solution. Nine to eleven different hot spots in both cancerous and non-cancerous regions in human breast cancers were captured per slide using *×*400 magnification for counting DLX5-positive and -negative spindle-shape fibroblast-like cells. For double fluorescence immunostaining, we stained human breast cancer tissues with anti-human DLX5 rabbit IgG and anti-human α-SMA or POSTN mouse IgG overnight at 4. Alexa Fluor 594-conjugated anti-rabbit IgG and Alexa Fluor 488-conjugated anti-mouse IgG were incubated for 1 h. The tissue was mounted with SlowFade™ Diamond Antifade Mountant with DAPI. Images were acquired with an Axioplan 2 microscope (Carl Zeiss) and prepared using ZEN Pro (Carl Zeiss). Green and red pseudo colors were assigned to DLX5 and α-SMA or POSTN for visibility, respectively.

### Measurement of TGF-β activity

To collect C.M., 0.4×10^5^ 218TGpp CSIV-GFP or FLAG-DLX5 cells were seeded in a 6-well plate. Those cells were treated with DOX for 48 h (0.5 µg/ml) before the medium was replaced with 1 mL of DMEM supplemented with FBS (10%) and DOX and cultured for an additional 48 h. For preparation of C.M. of MCF7-ras, 8.0×10^5^ cells were seeded and cultured for 72 h. The medium was replaced with 6 ml of culture medium for incubation for 48 h. C.M. was passed through a 0.45-µm-pore filter and stored at −80. TGF-β activity of C.M. was determined by a procedure described in our previous papers (Kojima *et al*., 2010; Mezawa *et al*., 2023). Briefly, mink lung epithelial cells (MLEC) stably harboring a TGF-β reporter construct which was constituted with TGF-β responsive element of the human *PAI-1* (*SERPINE1*) gene and firefly luciferase were treated with twofold diluted C.M. for 14 h (Abe *et al*, 1994). Cell lysate was prepared with passive lysis buffer (Promega, E194A) and luciferase activity was measured with Dual-Glo® Luciferase Assay System and GloMax® Discover System (Promega, E2940).

To evaluate TGF-β activity in the co-culture condition, 0.8×10^5^ of 218TGpp-CSIV-GFP or -FLAG-DLX5 were seeded on 6-well plate. Those cells were treated with DOX (0.5 µg/ml) on the next day for 4 days. 1.35 ×10^5^ of MLECs which had been transfected with Renilla luciferase encoding vector pRL-CMV (PicaGene, 307-05601) prior to 48 h were seeded on the fibroblasts treated with DOX for three days. The same number of MLECs were seeded on 4 cm dishes for monoculture. Following co-culture for 18 h, cell lysate was prepared with passive lysis buffer (Promega #E194A). Firefly luciferase activity was measured using the Bright-Glo™ Luciferase Assay System (Promega, E2620), whereas the Dual-Glo® Luciferase Assay System and GloMax® Discover System (Promega, E2940) were used to detect Renilla luciferase activity and normalize for the number of attached MLECs by calculating fLuc/rLuc ratios.

### Xenograft assay

The xenograft experiment was performed as described in a previous study (Matsumura *et al*., 2019). In brief, 1×10^5^ DCIStb cells and 3×10^5^ 218TGpp-pCDH FLAG-DLX5 IRES NEO or 218TGpp-pCDH mVenus IRES NEO cells were suspended in 50% Matrigel and injected subcutaneously into NOD/Shi-scid,IL-2RγKO (NOG) mice. After 29 days, tumors were resected and subjected to fixation and histological analysis. Budding-like regions were quantified by the following procedure: a region containing at least 10 ball-like cancer cell clusters, each composed of 3–20 cells, within a single field of view (1.2×2.4 mm) was identified, and the surrounding area enriched for similar cell clusters was manually delineated using SLIDEVIEW VS200 (Evident Scientific).

### Statistical analysis and reproducibility

Tests of statistical significance are described in each figure legend and in the Methods section. *P* < 0.05 is considered to indicate a statistically significant difference. The number of replicates is also described in each figure legend.

## Data availability

The source of the original sequence data was written above. The integrated scRNA-seq data presented in this study will be shared upon reasonable request. Bulk RNA-seq and CUT&RUN data will be publicly available in the Gene Expression Omnibus upon manuscript acceptance.

## Ethics statement

All scRNA-seq data from human specimens used in the meta-analyses were obtained from public databases. The use of formalin-fixed, paraffin-embedded human breast cancer tissue specimens for immunostaining in this study was reviewed and approved by the Juntendo University Ethics Review Board (approval number: H16-0160). The animal experiments were approved by the Animal Research Ethics Committee of the Juntendo University Faculty of Medicine (approval number: 1035).

## Author contributions

Y.M. and A.O. contributed to the concept, study design and interpretation of the data. Y.M., K.M. and K.H. contributed to investigation. Y.M. performed formal analysis of scRNA-seq, bulk RNA-seq, CUT&RUN and all wet experiments. K.K., L.Y. and R.M. contributed to the data acquisition and formal analysis for scATAC-seq. Y.M., K.M., K.Y., T.S., H.O., R.S. and G.K. contributed to method and material development. Y.M. and A.O. wrote and edited the manuscript. All authors read and approved the final manuscript.

## Acknowledgements

We thank Dr. Hiromichi Tsurui at Juntendo University for supporting bioinformatic analysis and Drs. Harumi Saeki and Kazunori Kajino at Juntendo University for supporting pathological analysis. We thank Division of Cell Biology, the Laboratory of Molecular and Biochemical Research, and Atopy Research Center at Juntendo University for technical assistance. We thank Ms. Yasuno Sugihara, Ms. Chihiro Yatomi, Mr. Yuta Imamura, and Mr. Ryotaro Yamamoto for their technical support on this project during Seminar in Basic Medicine at Juntendo University School of Medicine. This work was supported by Chugai Foundation for Innovative Drug Discovery Science, the Leading Center for Development and Research on Cancer Medicine at Juntendo University, Center of Genomic and Regeneration Medicine at Juntendo University, and Institute for Diseases of Old Age at Juntendo University. This work was supported by JSPS KAKENHI Grant Numbers 22K20837 and 25K18884 for Y.M. and 18K07207 for A.O.

---

**Figure EV1.**
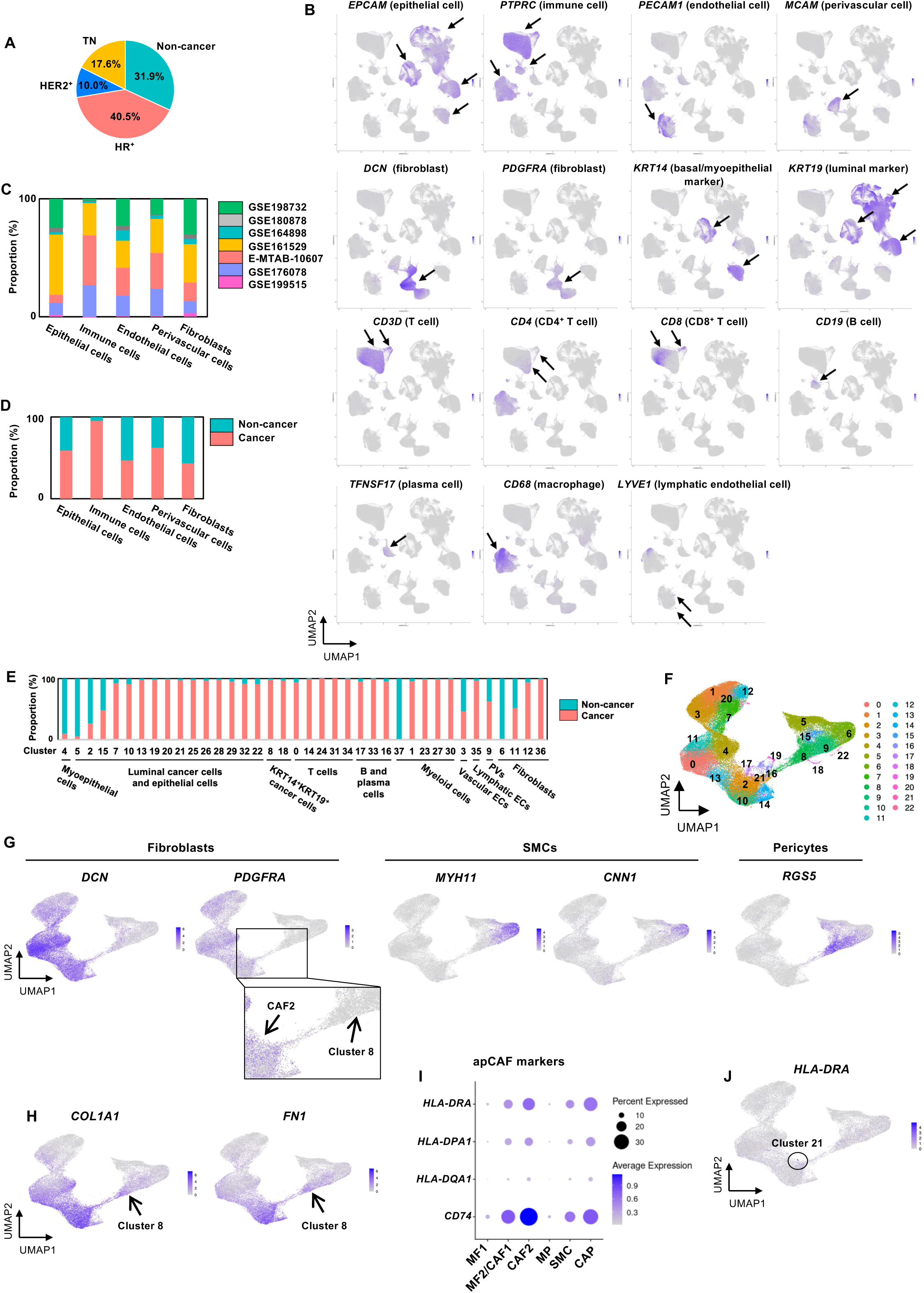
Cell type marker expression and information on patients and cell types. **(A)** Donor conditions were categorized as non-cancer or by intrinsic subtype of breast cancer. HR, hormone receptor-positive; TN, triple-negative; HER2, human epidermal growth factor receptor 2-positive. **(B)** Uniform manifold approximation and projection (UMAP) for cell type and subset markers. Arrows indicate regions strongly expressing indicated markers. **(C, D)** Proportions of source data and donor condition are displayed for each cell type. **(E)** Proportions of donor condition are displayed for each cell type or subset. **(F)** UMAP of fibroblasts and PVs (perivascular cells) before filtering, as described in Methods. **(G)** Gene expression of markers for fibroblasts (*DCN* and *PDGFRA*) and PVs (*MYH11*, *CNN1* and *RGS5*) is displayed on UMAP. **(H)** Gene expression is displayed on UMAP. Arrows, cells potentially undergoing pericyte-to-fibroblast transition. **(I)** Balloon plot of apCAF markers. **(J)** Expression level of *HLA-DRA* is displayed on UMAP. Circle, cluster 21 with the highest expression of *HLA-DRA*.

**Figure EV2.**
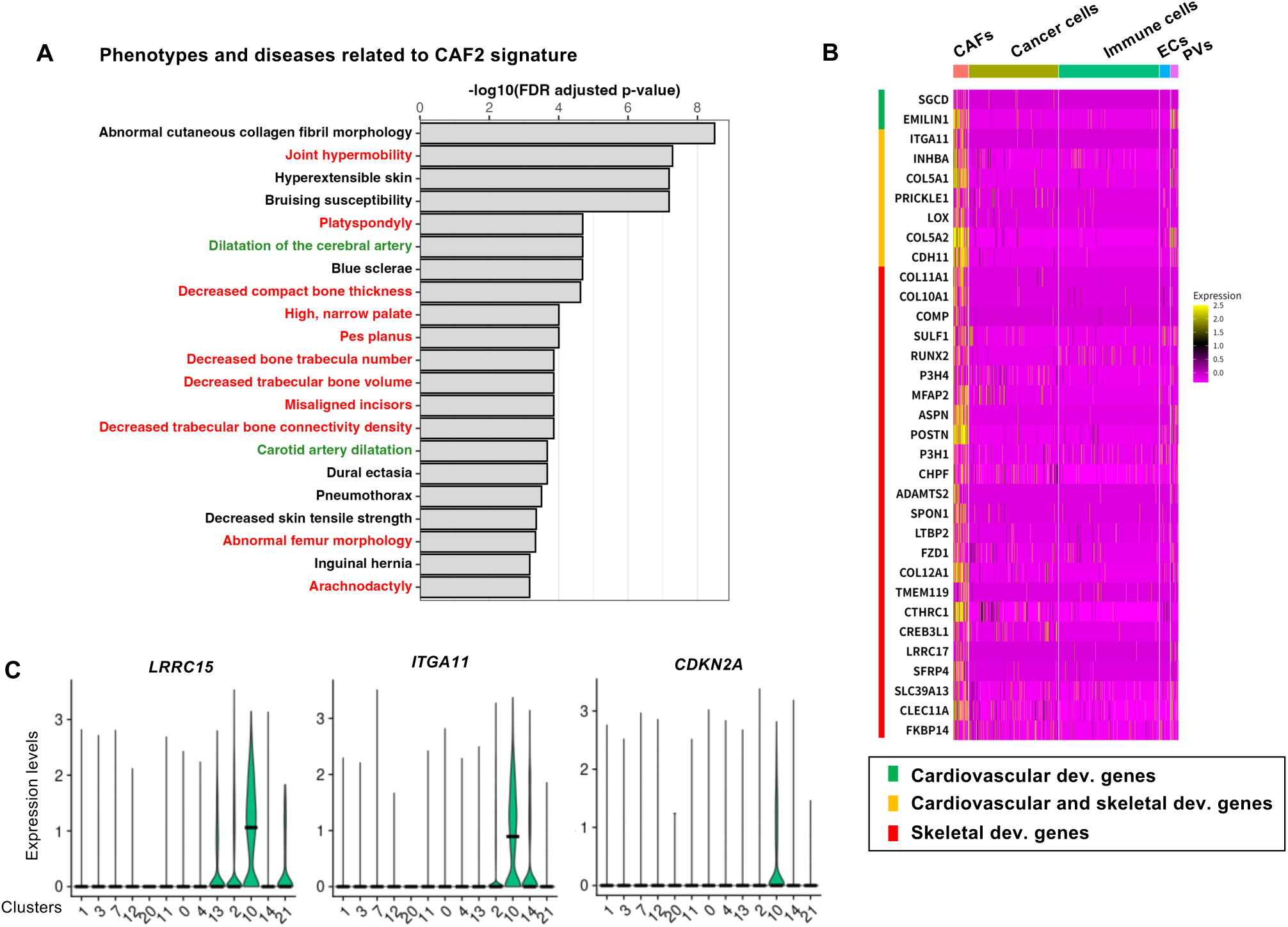
Characteristics of the CAF2 cluster. **(A)** Enrichment analysis for the Human Phenotype Ontology and the Mouse Genome Database by using PhenoExam (Cisterna *et al*., 2022). GO terms associated with the cardiovascular or skeletal system are shown in green and red, respectively. Terms with an FDR < 0.001 are shown. **(B)** Expression levels of genes highlighted with a gray background in Figure 2D are shown across major cell types in cancer tissues. These genes were selected based on their predominant expression in CAFs within cancer tissues. **(C)** Violin plots for indicated genes in the current integrated data set are shown for each fibroblast cluster.

**Figure EV3.**
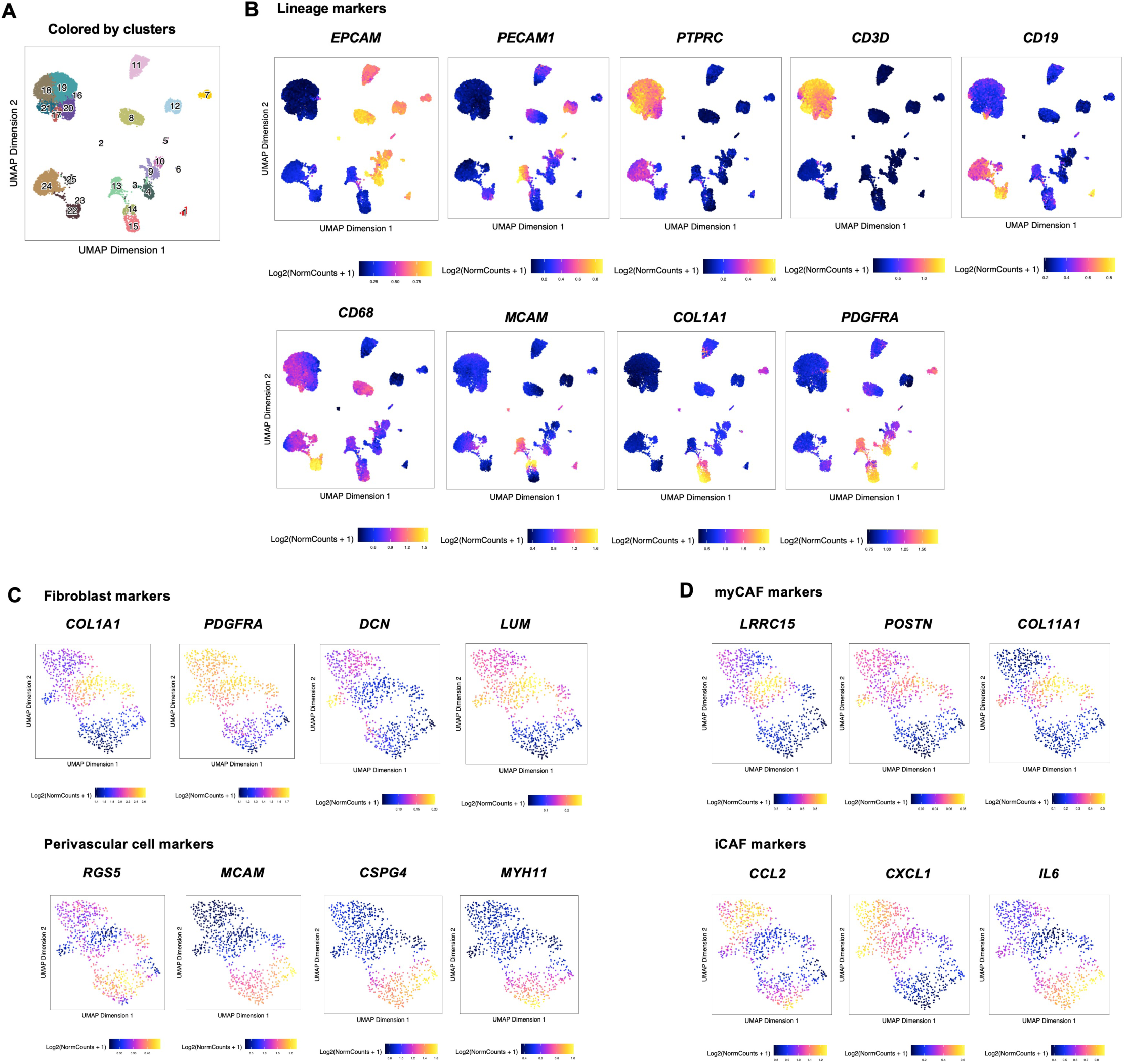
Data related to scATAC-seq for human breast cancer patients. **(A)** UMAP plot of all cells that passed quality control. Numbers indicate clusters. **(B)** Gene scores (GSs) for major lineage markers are shown. **(C, D)** GSs for the indicated genes are shown.

**Figure EV4.**
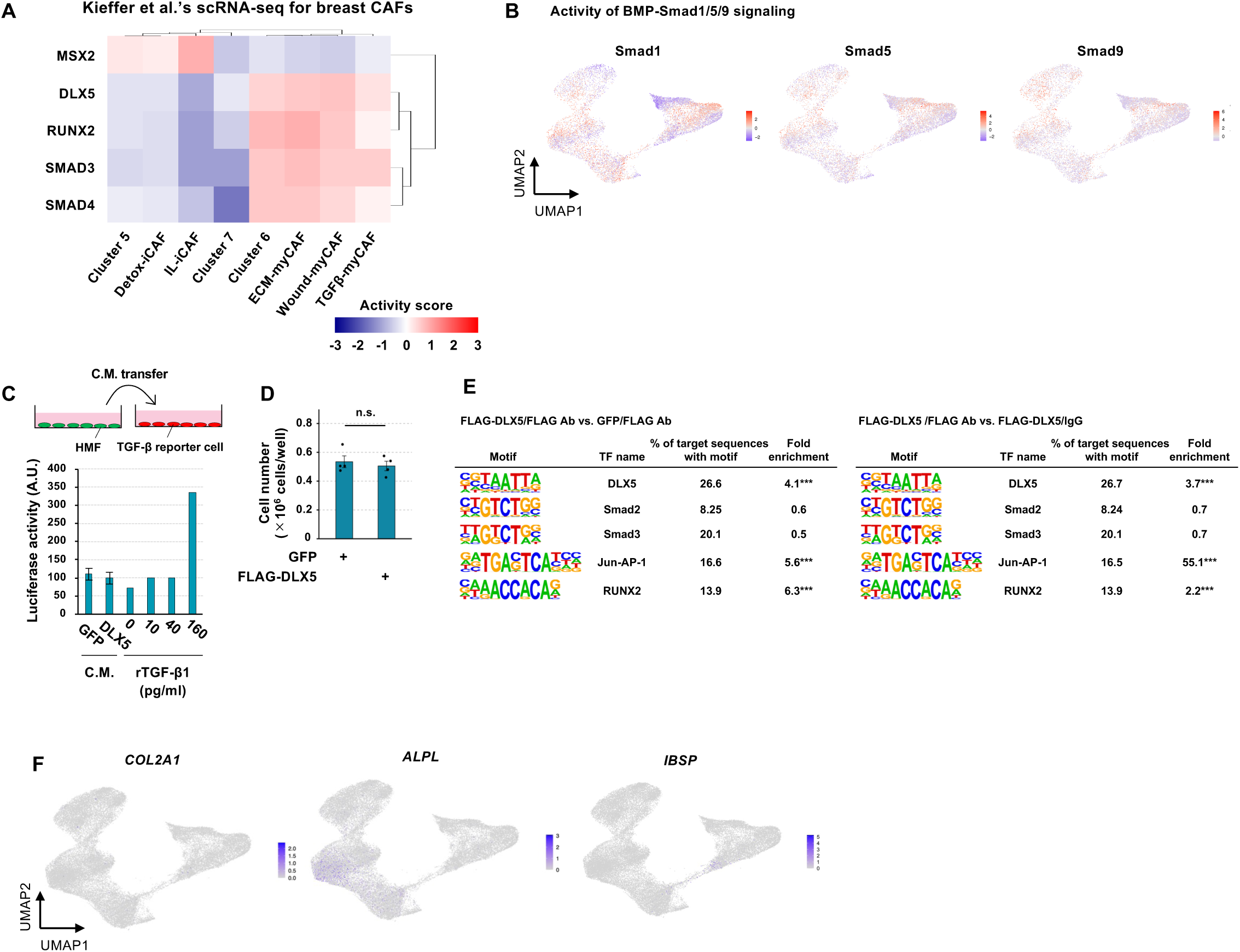
Characteristics of the CAF2 cluster and additional data on DLX5 function in TGF-β activity. **(A)** Transcription factor activities were estimated using scRNA-seq data for the CAF-S1 subset. **(B)** Activities of Smads involved in the BMP-Smad1/5/9 signaling pathway were estimated using the current integrated scRNA-seq dataset. **(C)** TGF-β activity in conditioned medium (C.M.) derived from GFP- or DLX5-HMFs was assayed using mink lung epithelial cells stably harboring a TGF-β reporter construct (n = 4). **(D)** GFP- or DLX5-HMFs were seeded on 6-well plate and cultured with the standard condition supplemented with DOX (0.5 µg/ml) for 4 days and cell number was measured. n.s., not significant by Student’s t-test (n = 4). Error bars, S.E. **(E)** Motif enrichment analysis of the FLAG-DLX5/FLAG Ab sample compared with the FLAG-DLX5/IgG or GFP/FLAG Ab samples. *p*-values were calculated using the hypergeometric test and adjusted for multiple testing using Benjamini–Hochberg method in HOMER (\*\*\**q* < 0.001). **(F)** Gene expression is displayed on UMAP.

**Figure EV5.**
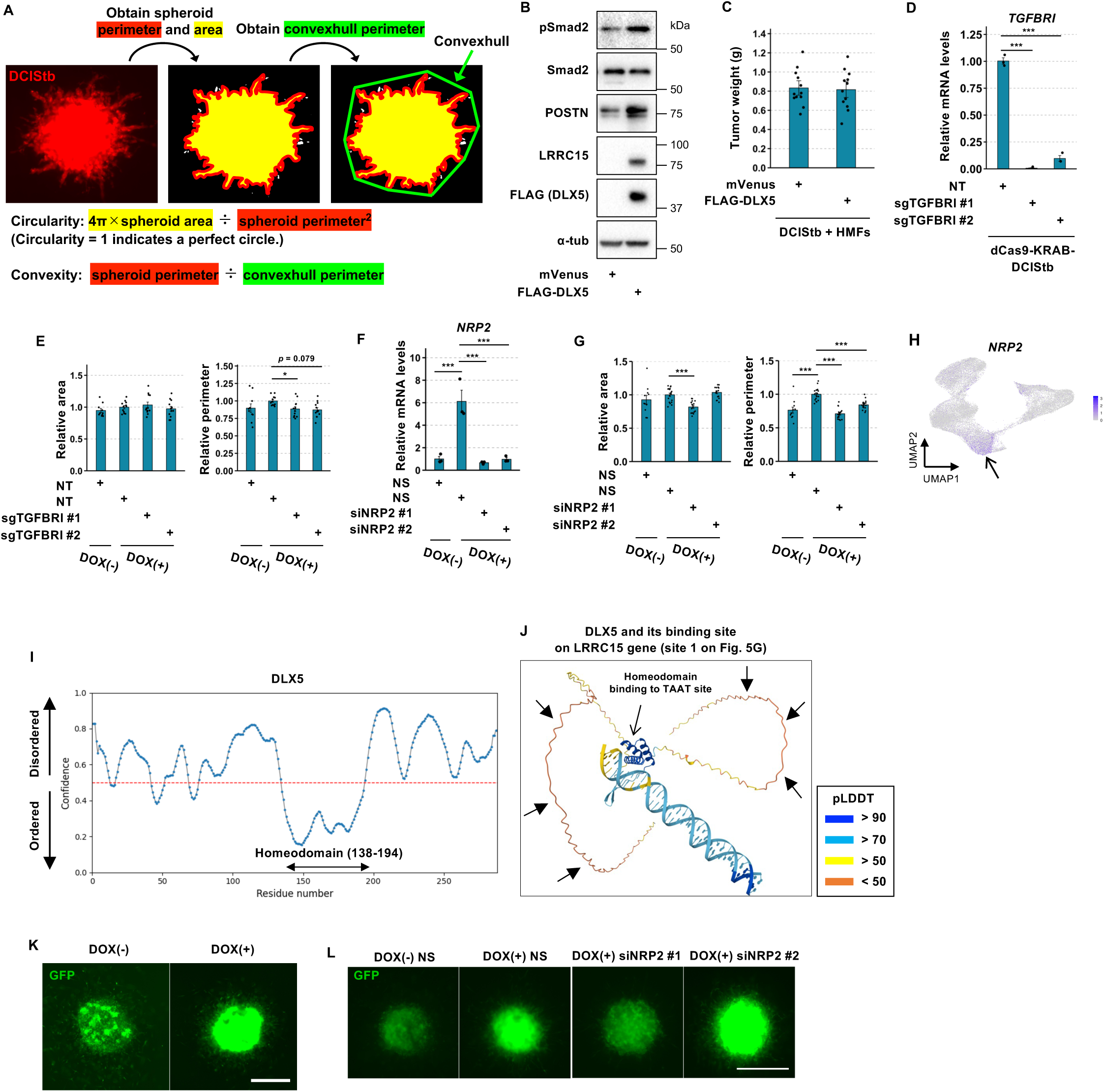
Additional data on phenotypic characterization of DLX5-HMFs. **(A)** Image analysis workflow for evaluating cancer cell invasion in the 3D co-culture assay using ImageJ. **(B)** HMFs constitutively expressing mVenus or FLAG-DLX5 with pCDH lentiviral vectors were subjected to western blot analysis. **(C)** These cells were co-injected with DCIStb cells into NOG mice, and tumors that developed by day 29 were harvested and weighed (n = 12). **(D)** Confirmation of *TGFBRI* knockdown in DCIStb cells expressing dCas9-KRAB and sgTGFBRI. NT, non-targeting sgRNA. \*\*\**p* < 0.001 by Dunnett’s t-test (n = 3). **(E)** The area and perimeter of tdTomato-positive DCIStb cells in the experiments shown in Fig. 6I and J are shown. \**p* < 0.05 by Dunnett’s t-test (n = 10 to 12 from three independent experiments). **(F)** FLAG-DLX5-HMFs transfected with siNRP2 or non-silencing siRNA (NS) were treated with DOX (0.5 μg/mL) for 48 h and subjected to real-time PCR. **(G)** The area and perimeter of tdTomato-positive DCIStb cells in the experiments shown in Fig. 6L and M are shown. \*\*\**p* < 0.001 by Dunnett’s t-test (n = 12 to 18 from three independent experiments). **(H)** *NRP2* expression is shown on a UMAP of human breast cancer scRNA-seq data. Arrow, the CAF2 cluster showing high *NRP2* expression. **(I)** Intrinsically disordered regions (IDRs) of human DLX5 (NP_005212.1) predicted by the PrDOS tool with the default settings. **(J)** Structure of human DLX5 (NP_005212.1) bound to the indicated DNA, as predicted by AlphaFold3. pLDDT, predicted local distance difference test. Arrows indicate putative IDRs in the N- and C-terminal regions, showing low pLDDT. **(K, L)** GFP fluorescence derived from DLX5-HMFs in spheroids (Figure 6B and 6L) is shown. Scale bars, 500 µm.

## References

Abe M, Harpel JG, Metz CN, Nunes I, Loskutoff DJ, Rifkin DB (1994) An assay for transforming growth factor-beta using cells transfected with a plasminogen activator inhibitor-1 promoter-luciferase construct. Anal Biochem 216: 276–284

Acampora D, Merlo GR, Paleari L, Zerega B, Postiglione MP, Mantero S, Bober E, Barbieri O, Simeone A, Levi G (1999) Craniofacial, vestibular and bone defects in mice lacking the Distal-less-related gene Dlx5. Development 126: 3795–3809

Amarasekara DS, Kim S, Rho J (2021) Regulation of Osteoblast Differentiation by Cytokine Networks. Int J Mol Sci 22: 2851

Bartoschek M, Oskolkov N, Bocci M, Lovrot J, Larsson C, Sommarin M, Madsen CD, Lindgren D, Pekar G, Karlsson G et al (2018) Spatially and functionally distinct subclasses of breast cancer-associated fibroblasts revealed by single cell RNA sequencing. Nat Commun 9: 5150

Bhat-Nakshatri P, Gao H, Sheng L, McGuire PC, Xuei X, Wan J, Liu Y, Althouse SK, Colter A, Sandusky G et al (2021) A single-cell atlas of the healthy breast tissues reveals clinically relevant clusters of breast epithelial cells. Cell Rep Med 2: 100219

Cao J, Spielmann M, Qiu X, Huang X, Ibrahim DM, Hill AJ, Zhang F, Mundlos S, Christiansen L, Steemers FJ et al (2019) The single-cell transcriptional landscape of mammalian organogenesis. Nature 566: 496–502

Chong S, Dugast-Darzacq C, Liu Z, Dong P, Dailey GM, Cattoglio C, Heckert A, Banala S, Lavis L, Darzacq X et al (2018) Imaging dynamic and selective low-complexity domain interactions that control gene transcription. Science 361: eaar2555

Cisterna A, Gonzalez-Vidal A, Ruiz D, Ortiz J, Gomez-Pascual A, Chen Z, Nalls M, Faghri F, Hardy J, Diez I et al (2022) PhenoExam: gene set analyses through integration of different phenotype databases. BMC Bioinformatics 23: 567

Croizer H, Mhaidly R, Kieffer Y, Gentric G, Djerroudi L, Leclere R, Pelon F, Robley C, Bohec M, Meng A et al (2024) Deciphering the spatial landscape and plasticity of immunosuppressive fibroblasts in breast cancer. Nat Commun 15: 2806

David CJ, Massague J (2018) Contextual determinants of TGFbeta action in development, immunity and cancer. Nat Rev Mol Cell Biol 19: 419–435

Delgado AP, Nemajerova A, Nelson BJ, Rao M, Li J, Marchenko N, Preall J, Moll UM, Egeblad M, Powers S (2024) Single-cell transcriptome analysis reveals evolutionarily conserved features during the transition from normal breast stromal cells to cancer-associated fibroblasts. bioRxiv 10.1101/2022.05.05.490693 [PREPRINT]

Depew MJ, Liu JK, Long JE, Presley R, Meneses JJ, Pedersen RA, Rubenstein JL (1999) Dlx5 regulates regional development of the branchial arches and sensory capsules. Development 126: 3831–3846

Dominguez CX, Muller S, Keerthivasan S, Koeppen H, Hung J, Gierke S, Breart B, Foreman O, Bainbridge TW, Castiglioni A et al (2019) Single-cell RNA sequencing reveals stromal evolution into LRRC15+ myofibroblasts as a determinant of patient response to cancer immunotherapy. Cancer Discov 10: 232–253

Elyada E, Bolisetty M, Laise P, Flynn WF, Courtois ET, Burkhart RA, Teinor JA, Belleau P, Biffi G, Lucito MS et al (2019) Cross-Species Single-Cell Analysis of Pancreatic Ductal Adenocarcinoma Reveals Antigen-Presenting Cancer-Associated Fibroblasts. Cancer Discov 9: 1102–1123

Gaggioli C, Hooper S, Hidalgo-Carcedo C, Grosse R, Marshall JF, Harrington K, Sahai E (2007) Fibroblast-led collective invasion of carcinoma cells with differing roles for RhoGTPases in leading and following cells. Nat Cell Biol 9: 1392–1400

Gao Y, Li J, Cheng W, Diao T, Liu H, Bo Y, Liu C, Zhou W, Chen M, Zhang Y et al (2024) Cross-tissue human fibroblast atlas reveals myofibroblast subtypes with distinct roles in immune modulation. Cancer Cell 42: 1764–1783

Granja JM, Corces MR, Pierce SE, Bagdatli ST, Choudhry H, Chang HY, Greenleaf WJ (2021) ArchR is a scalable software package for integrative single-cell chromatin accessibility analysis. Nat Genet 53: 403–411

Gray GK, Li CM, Rosenbluth JM, Selfors LM, Girnius N, Lin JR, Schackmann RCJ, Goh WL, Moore K, Shapiro HK et al (2022) A human breast atlas integrating single-cell proteomics and transcriptomics. Dev Cell 57: 1400–1420

Hao Y, Stuart T, Kowalski MH, Choudhary S, Hoffman P, Hartman A, Srivastava A, Molla G, Madad S, Fernandez-Granda C et al (2024) Dictionary learning for integrative, multimodal and scalable single-cell analysis. Nat Biotechnol 42: 293–304

Hosaka K, Yang Y, Seki T, Fischer C, Dubey O, Fredlund E, Hartman J, Religa P, Morikawa H, Ishii Y et al (2016) Pericyte-fibroblast transition promotes tumor growth and metastasis. Proc Natl Acad Sci U S A 113: E5618–5627

Hu M, Yao J, Carroll DK, Weremowicz S, Chen H, Carrasco D, Richardson A, Violette S, Nikolskaya T, Nikolsky Y et al (2008) Regulation of in situ to invasive breast carcinoma transition. Cancer Cell 13: 394–406

Islam R, Mishra J, Bodas S, Bhattacharya S, Batra SK, Dutta S, Datta K (2022) Role of Neuropilin-2-mediated signaling axis in cancer progression and therapy resistance. Cancer Metastasis Rev 41: 771–787

Kieffer Y, Hocine HR, Gentric G, Pelon F, Bernard C, Bourachot B, Lameiras S, Albergante L, Bonneau C, Guyard A et al (2020) Single-Cell Analysis Reveals Fibroblast Clusters Linked to Immunotherapy Resistance in Cancer. Cancer Discov 10: 1330–1351

Kojima Y, Acar A, Eaton EN, Mellody KT, Scheel C, Ben-Porath I, Onder TT, Wang ZC, Richardson AL, Weinberg RA et al (2010) Autocrine TGF-beta and stromal cell-derived factor-1 (SDF-1) signaling drives the evolution of tumor-promoting mammary stromal myofibroblasts. Proc Natl Acad Sci U S A 107: 20009–20014

Komori T, Yagi H, Nomura S, Yamaguchi A, Sasaki K, Deguchi K, Shimizu Y, Bronson RT, Gao YH, Inada M et al (1997) Targeted disruption of Cbfa1 results in a complete lack of bone formation owing to maturational arrest of osteoblasts. Cell 89: 755–764

Korsunsky I, Millard N, Fan J, Slowikowski K, Zhang F, Wei K, Baglaenko Y, Brenner M, Loh PR, Raychaudhuri S (2019) Fast, sensitive and accurate integration of single-cell data with Harmony. Nat Methods 16: 1289–1296

Krishnamurty AT, Shyer JA, Thai M, Gandham V, Buechler MB, Yang YA, Pradhan RN, Wang AW, Sanchez PL, Qu Y et al (2022) LRRC15(+) myofibroblasts dictate the stromal setpoint to suppress tumour immunity. Nature 611: 148–154

Kumar T, Nee K, Wei R, He S, Nguyen QH, Bai S, Blake K, Pein M, Gong Y, Sei E et al (2023) A spatially resolved single-cell genomic atlas of the adult human breast. Nature 620: 181–191

Kumegawa K, Takahashi Y, Saeki S, Yang L, Nakadai T, Osako T, Mori S, Noda T, Ohno S, Ueno T et al (2022) GRHL2 motif is associated with intratumor heterogeneity of cis-regulatory elements in luminal breast cancer. NPJ Breast Cancer 8: 70

Lavie D, Ben-Shmuel A, Erez N, Scherz-Shouval R (2022) Cancer-associated fibroblasts in the single-cell era. Nat Cancer 3: 793–807

Lendahl U, Muhl L, Betsholtz C (2022) Identification, discrimination and heterogeneity of fibroblasts. Nat Commun 13: 3409

Levi G, Narboux-Neme N, Cohen-Solal M (2022) DLX Genes in the Development and Maintenance of the Vertebrate Skeleton: Implications for Human Pathologies. Cells 11: 3277

Li J, Wang W, Li S, Qiao Z, Jiang H, Chang X, Zhu Y, Tan H, Ma X, Dong Y et al (2024) Smad2/3/4 complex could undergo liquid liquid phase separation and induce apoptosis through TAT in hepatocellular carcinoma. Cancer Cell Int 24: 176

Liu Y, Chen X, Dai Y, Jia Y, Kieffer Y, Xie L, Zhou Z, Tyler L, Kim AC, Biffi G et al (2026) Molecular phenotypes and spatial archetypes: A new framework for cancer-associated fibroblasts. Cancer Cell 44: 1346–1367

Maira M, Long JE, Lee AY, Rubenstein JL, Stifani S (2010) Role for TGF-beta superfamily signaling in telencephalic GABAergic neuron development. J Neurodev Disord 2: 48–60

Mariathasan S, Turley SJ, Nickles D, Castiglioni A, Yuen K, Wang Y, Kadel EE, III, Koeppen H, Astarita JL, Cubas R et al (2018) TGFbeta attenuates tumour response to PD-L1 blockade by contributing to exclusion of T cells. Nature 554: 544–548

Massague J, Sheppard D (2023) TGF-beta signaling in health and disease. Cell 186: 4007–4037

Matsumura Y, Ito Y, Mezawa Y, Sulidan K, Daigo Y, Hiraga T, Mogushi K, Wali N, Suzuki H, Itoh T et al (2019) Stromal fibroblasts induce metastatic tumor cell clusters via epithelial-mesenchymal plasticity. Life Sci Alliance 2: e201900425

McGinnis CS, Murrow LM, Gartner ZJ (2019) DoubletFinder: Doublet Detection in Single-Cell RNA Sequencing Data Using Artificial Nearest Neighbors. Cell Syst 8: 329–337

Mezawa Y, Daigo Y, Takano A, Miyagi Y, Yokose T, Yamashita T, Morimoto C, Hino O, Orimo A (2019) CD26 expression is attenuated by TGF-beta and SDF-1 autocrine signaling on stromal myofibroblasts in human breast cancers. Cancer Med 8: 3936–3948

Mezawa Y, Orimo A (2022) Phenotypic heterogeneity, stability and plasticity in tumor-promoting carcinoma-associated fibroblasts. FEBS J 289: 2429–2447

Mezawa Y, Wang T, Daigo Y, Takano A, Miyagi Y, Yokose T, Yamashita T, Yang L, Maruyama R, Seimiya H et al (2023) Glutamine deficiency drives transforming growth factor-beta signaling activation that gives rise to myofibroblastic carcinoma-associated fibroblasts. Cancer Sci 114: 4376–4387

Miyazawa K, Itoh Y, Fu H, Miyazono K (2024) Receptor-activated transcription factors and beyond: multiple modes of Smad2/3-dependent transmission of TGF-beta signaling. J Biol Chem 300: 107256

Morini M, Astigiano S, Gitton Y, Emionite L, Mirisola V, Levi G, Barbieri O (2010) Mutually exclusive expression of DLX2 and DLX5/6 is associated with the metastatic potential of the human breast cancer cell line MDA-MB-231. BMC Cancer 10: 649

Muir AM, Ren Y, Butz DH, Davis NA, Blank RD, Birk DE, Lee SJ, Rowe D, Feng JQ, Greenspan DS (2014) Induced ablation of Bmp1 and Tll1 produces osteogenesis imperfecta in mice. Hum Mol Genet 23: 3085–3101

Murrow LM, Weber RJ, Caruso JA, McGinnis CS, Phong K, Gascard P, Rabadam G, Borowsky AD, Desai TA, Thomson M et al (2022) Mapping hormone-regulated cell-cell interaction networks in the human breast at single-cell resolution. Cell Syst 13: 644–664

Nixon BG, Gao S, Wang X, Li MO (2023) TGFbeta control of immune responses in cancer: a holistic immuno-oncology perspective. Nat Rev Immunol 23: 346–362

Okubo S, Mezawa Y, Wang Z, Acar A, Ito Y, Takano A, Miyagi Y, Yokose T, Yamashita T, Daigo Y et al (2025) Endoglin mediates the tumor- and metastasis-promoting traits of stromal myofibroblasts in human breast carcinomas. Mol Oncol 19: 2557–2573

Pal B, Chen Y, Vaillant F, Capaldo BD, Joyce R, Song X, Bryant VL, Penington JS, Di Stefano L, Tubau Ribera N et al (2021) A single-cell RNA expression atlas of normal, preneoplastic and tumorigenic states in the human breast. EMBO J 40: e107333

Polanska UM, Mellody KT, Orimo A (2010) Tumour-Promoting Stromal Myofibroblasts in Human Carcinomas RG Bagley (ed), The Tumor Microenvironment, Cancer Drug Discovery and Development: Chapter 16, 325–349

Purcell JW, Tanlimco SG, Hickson J, Fox M, Sho M, Durkin L, Uziel T, Powers R, Foster K, McGonigal T et al (2018) LRRC15 Is a Novel Mesenchymal Protein and Stromal Target for Antibody-Drug Conjugates. Cancer Res 78: 4059–4072

Richards AJ, Yates JR, Williams R, Payne SJ, Pope FM, Scott JD, Snead MP (1996) A family with Stickler syndrome type 2 has a mutation in the COL11A1 gene resulting in the substitution of glycine 97 by valine in alpha 1 (XI) collagen. Hum Mol Genet 5: 1339–1343

Serini G, Gabbiani G (1999) Mechanisms of myofibroblast activity and phenotypic modulation. Exp Cell Res 250: 273–283

Shen G (2005) The role of type X collagen in facilitating and regulating endochondral ossification of articular cartilage. Orthod Craniofac Res 8: 11–17

Shirakabe K, Terasawa K, Miyama K, Shibuya H, Nishida E (2001) Regulation of the activity of the transcription factor Runx2 by two homeobox proteins, Msx2 and Dlx5. Genes Cells 6: 851–856

Tan Y, Timakhov RA, Rao M, Altomare DA, Xu J, Liu Z, Gao Q, Jhanwar SC, Di Cristofano A, Wiest DL et al (2008) A novel recurrent chromosomal inversion implicates the homeobox gene Dlx5 in T-cell lymphomas from Lck-Akt2 transgenic mice. Cancer Res 68: 1296–1302

Tietscher S, Wagner J, Anzeneder T, Langwieder C, Rees M, Sobottka B, de Souza N, Bodenmiller B (2023) A comprehensive single-cell map of T cell exhaustion-associated immune environments in human breast cancer. Nat Commun 14: 98

Tirosh I, Suva ML (2024) Cancer cell states: Lessons from ten years of single-cell RNA-sequencing of human tumors. Cancer Cell 42: 1497–1506

Wang H, Li N, Liu Q, Guo J, Pan Q, Cheng B, Xu J, Dong B, Yang G, Yang B et al (2023) Antiandrogen treatment induces stromal cell reprogramming to promote castration resistance in prostate cancer. Cancer Cell 41: 1345–1362

Wishman MD, Sgrignoli WM, Patterson BM, Nepola JV, Wolf BR, Bozoghlian M, Lane CM, Coleman MC, Galvin JW (2025) A review of periostin in orthopedics. Osteoarthr Cartil Open 7: 100600

Wittmann P, Grubinger M, Groger C, Huber H, Sieghart W, Peck-Radosavljevic M, Mikulits W (2015) Neuropilin-2 induced by transforming growth factor-beta augments migration of hepatocellular carcinoma cells. BMC Cancer 15: 909

Wu SZ, Al-Eryani G, Roden DL, Junankar S, Harvey K, Andersson A, Thennavan A, Wang C, Torpy JR, Bartonicek N et al (2021) A single-cell and spatially resolved atlas of human breast cancers. Nat Genet 53: 1334–1347

Ye J, Baer JM, Faget DV, Morikis VA, Ren Q, Melam A, Delgado AP, Luo X, Bagchi SM, Belle JI et al (2024) Senescent CAFs Mediate Immunosuppression and Drive Breast Cancer Progression. Cancer Discov 14: 1302–1323

Zeisberg EM, Potenta S, Xie L, Zeisberg M, Kalluri R (2007) Discovery of endothelial to mesenchymal transition as a source for carcinoma-associated fibroblasts. Cancer Res 67: 10123–10128

Zerucha T, Stuhmer T, Hatch G, Park BK, Long Q, Yu G, Gambarotta A, Schultz JR, Rubenstein JL, Ekker M (2000) A highly conserved enhancer in the Dlx5/Dlx6 intergenic region is the site of cross-regulatory interactions between Dlx genes in the embryonic forebrain. J Neurosci 20: 709–721

Zheng H, An M, Luo Y, Diao X, Zhong W, Pang M, Lin Y, Chen J, Li Y, Kong Y et al (2024) PDGFRalpha(+)ITGA11(+) fibroblasts foster early-stage cancer lymphovascular invasion and lymphatic metastasis via ITGA11-SELE interplay. Cancer Cell 42: 682–700

